# Selection of Ethanol Tolerant Strains of *Candida albicans* by Repeated Ethanol Exposure Results in Strains with Reduced Susceptibility to Fluconazole

**DOI:** 10.1101/2023.09.13.557677

**Authors:** Andrew W. Day, Carol A. Kumamoto

## Abstract

*Candida albicans* is a commensal yeast that has important impacts on host metabolism and immune function, and can establish life-threatening infections in immunocompromised individuals. Previously, *C. albicans* colonization has been shown to contribute to the progression and severity of alcoholic liver disease. However, relatively little is known about how *C. albicans* responds to changing environmental conditions in the GI tract of individuals with alcohol use disorder, namely repeated exposure to ethanol. In this study, we repeatedly exposed *C. albicans* to high concentrations (10% vol/vol) of ethanol—a concentration that can be observed in the upper GI tract of humans following consumption of alcohol. Following this repeated exposure protocol, ethanol small colony (Esc) variants of *C. albicans* isolated from these populations exhibited increased ethanol tolerance, altered transcriptional responses to ethanol, and cross-resistance/tolerance to the frontline antifungal fluconazole. These Esc strains exhibited chromosomal copy number variations and carried polymorphisms in genes previously associated with the acquisition of fluconazole resistance during human infection. This study identifies a selective pressure that can result in evolution of fluconazole tolerance and resistance without previous exposure to the drug.

## Introduction

*Candida albicans* is an opportunistic pathogen causing severe morbidity and mortality in immunocompromised patients. *Candida spp* are the second most common cause of nosocomial bloodstream infection in the US, and cause about 70% of all fungal infections [1,2]. The most common species causing these fungal infections is *C. albicans*, the causative agent of oral thrush, vulvovaginitis, and bloodstream infections which result in mortality rates in immunocompromised patients ranging from roughly 30-56% [3–5]. Before *C. albicans* causes deadly bloodstream infections, it establishes commensalism within the gastrointestinal tract of humans [6,7]. Importantly, rates of antifungal resistance among *C. albicans* isolates have been rising throughout recent years [8], contributing to difficulty in treating bloodstream infections. Therefore, it is important to investigate factors that promote antifungal resistance whether acquired during infection or the preceding commensal state.

*C. albicans* commensalism has been studied particularly in disease states which predispose individuals to bloodstream infections. However, the environmental conditions in the GI tract and how they contribute to fungal physiology have been less studied. One condition of interest is the GI tract of individuals with alcohol use disorder (AUD). Recently, changes in the gut microbiome of individuals with alcohol use disorder, including a loss of fungal diversity, and blooms of *C. albicans* have been noted [9–11]. *C. albicans* blooms in the alcoholic GI tract promote the progression and increase the severity of alcoholic liver disease. Anti-*Saccharomyces cerevisiae* IgG antibodies (ASCA), which are thought to be broadly induced by fungi, show an inverse correlation with 6-month and 5-year survival rates in patients with alcoholic cirrhosis [9,10]. These findings suggest that fungal colonization can exacerbate diseases associated with alcohol use and abuse. However, there are still many questions surrounding why *C. albicans* relative abundance increases in the alcoholic GI tract and how *C. albicans* responds to environmental changes in the alcoholic GI tract.

One of the most intuitive features of the environment in the alcoholic GI tract is the increased presence of ethanol that members of the microbiota from the mouth through the proximal small intestine would be exposed to during bouts of consumption. Following consumption of one to two standard drinks of ethanol, the percentage of alcohol in the stomach and the small intestine can reach concentrations of roughly 10% ethanol [12,13]. To test how *C. albicans* responds to conditions that mimic this environment, we repeatedly cultured C. albicans in media with increasing amounts of ethanol, up to 10%. Model organisms such as *Escherichia coli* and *Saccharomyces cerevisiae* have been shown to evolve increased ethanol tolerance as a result of repeated exposure to ethanol [14,15]. We therefore hypothesized that *C. albicans* would evolve increased ethanol tolerance following repeated exposure to ethanol.

In this study, we find that populations of *C. albicans* cells repeatedly exposed to ethanol evolve increased ethanol tolerance. One ethanol tolerant strain showed increased ergosterol content during exposure to ethanol. We show that increased ergosterol content, whether induced by chemical stimulation or gain-of-function mutation, resulted in increased ethanol survival of *C. albicans*. Whole genome sequencing of strains with increased ethanol tolerance showed that these strains carried various chromosomal aneuploidies, some of which have been observed in fluconazole resistant and tolerant strains of *C. albicans*, and single nucleotide polymorphisms (SNPs). We also showed that the ethanol tolerant strains exhibited increased resistance and tolerance to azole drugs but not caspofungin or amphotericin B. These results suggest that *C. albicans* exposed to ethanol in the alcoholic GI tract could exhibit fluconazole resistance without previously being exposed to the antifungal drug.

## Results

### Selection of ethanol tolerant *C. albicans* isolates

The ability of *C. albicans* to evolve an altered response to ethanol exposure was investigated. To this end, *C. albicans* strain SC5314 was cultured eight times for 4 hrs each in tissue culture medium with 10% fetal bovine serum and 1X non-essential amino acids (referred to as medium A) with or without increasing concentrations of ethanol, as described in Materials and Methods (**Fig. 1A**). This process was repeated with five independent populations of cells.

**Figure 1.**
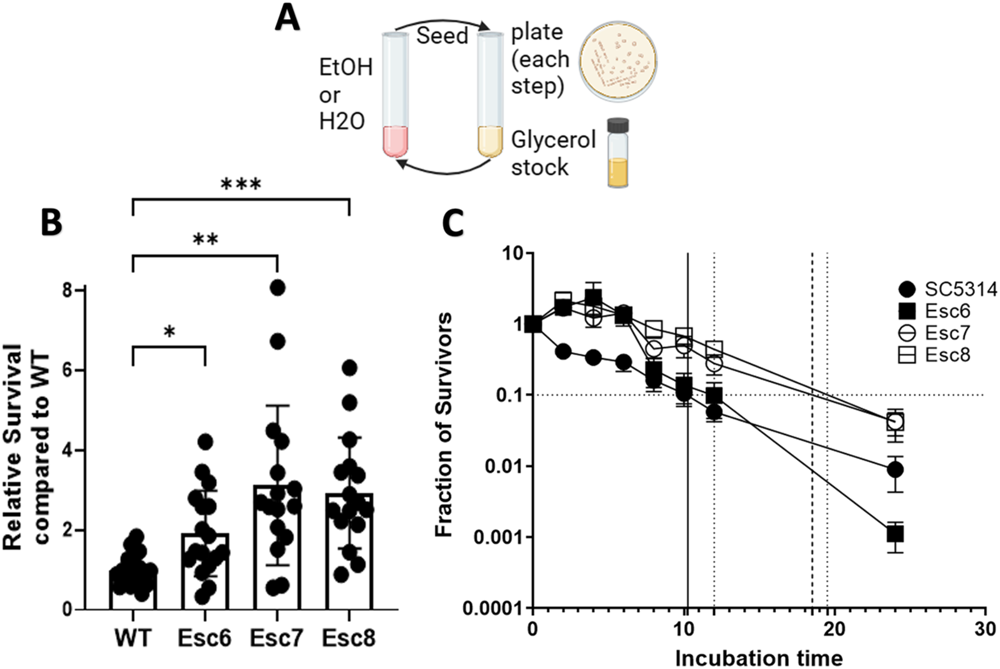
Isolation of Ethanol Tolerant Strains. **A)** The repeated ethanol exposure protocol is illustrated. 10^6^ cells of *C. albicans* strain SC5314 from 5 different biological replicate populations were inoculated into medium A with increasing concentrations of ethanol (1-10% vol/vol) or medium A diluted with water. Populations were exposed to 8 passages in ethanol or paired mock-exposed populations exposed to 8 passages in media with water. After each 4 hour ethanol or water exposure step, cells were seeded into YPD for recovery, and cultures were plated onto YPD-agar for examination of the populations. Recovery cultures were grown to post-exponential phase, and a glycerol stock was made. Cells from the recovery culture were then used for the next step. **B)** Relative survival of ethanol small colony (Esc) strains following a 4-hour ethanol exposure relative to WT ethanol survival was measured as described in Materials and Methods. Symbols show results from biological replicate cultures, bars show mean with standard deviation. Outliers were removed through ROUT, Q=1%, and statistical significance calculated by Brown-Forsythe and Welch ANOVA test. Each group represents 16-18 biological replicate cultures. (*, p < 0.0170; **, p < 0.0037; ***, p < 0.0004) **C)** Fraction of survivors was measured for 8-16 biological replicate cultures for each point. Timepoints taken were 0, 2, 4, 6, 8, 10, 12, or 24 hours of incubation time in 10% ethanol. Error bars show standard error for each timepoint. Horizontal line represents 90% death. Minimum duration of killing 90% (MDK90) values are plotted as vertical lines with solid black line representing WT and dashed lines representing Esc strains.

Throughout the repeated-ethanol exposure protocol, populations from each passage were plated onto YPD agar to monitor population dynamics. An increased number of small-sized colonies (50% of the size of normal sized colonies or less) was observed with greater numbers of passages and higher concentrations of ethanol (**Supplemental Fig. S1**). Small-sized colonies were rare in populations not exposed to ethanol. From these populations, small and normal sized colonies were isolated to determine their properties.

Normal-sized colonies were isolated from four different populations exposed repeatedly to ethanol, as well as from populations that were repeatedly exposed to water. Strains from normal-sized colonies were grown to post-exponential phase and added to medium A containing 10% ethanol or 10% water as described in Materials and Methods. Following the four-hour ethanol exposure at 37°C in a 5% CO_2_ atmosphere, cells were serially diluted, and plated to determine CFU and percentage survival. The final CFU of the 10% water and 10% ethanol cultures were counted and used to determine the final percentage survival of the ethanol cultures. Values were standardized within each experiment to the survival of SC5314 in 10% ethanol (**Supplemental Fig. S2**). Strains from normal-sized colonies from ethanol cultures showed no statistically significant increase in ethanol survival compared to SC5314 survival and were not studied further.

Ethanol small colony (Esc) variants were purified from different populations and tested as above to measure their ability to survive in ethanol. Relative survival values of SC5314, Esc6, Esc7, and Esc8 are shown in Fig. 1B. Esc6, Esc7, and Esc8 were isolated from different, independent populations and all showed significantly higher survival following the 4-hour ethanol exposure, compared to the parent strain. The growth rates of these strains were measured in YPD at 30⁰C; the strains showed increases in doubling times ranging from 1.37 to 2.12X greater than the doubling time of SC5314 shown in **Supplemental Fig. S3**.

The next experiment asked whether Esc strains had an increased ability to survive longer exposures to ethanol. SC5314, Esc6, Esc7, and Esc8 were grown overnight and then subcultured in medium A with 10% ethanol. CFU were determined for cultures at 0, 2, 4, 6, 8, 10, 12, and 24 hours of incubation. Fractions relative to the starting CFU of the cultures are shown in Fig. 1C. The results showed that Esc6, Esc7, and Esc8 outperformed WT *C. albicans* in ethanol survival from 0-6 hours and Esc7 and 8 were able to maintain higher ethanol survival relative to SC5314 through 24 hours. Esc6 had an early advantage in the first 6 hours compared to SC5314 but began to die rapidly after 12 hours. A common way to measure tolerance to antibiotics in bacterial systems is to determine the minimum duration of killing x fraction of the population (MDKx) [16]. We estimated the MDK90 for the strains by extrapolation and found that WT *C. albicans* had an MDK90 of 10 hrs, Esc6 had an MDK90 of 12 hrs, Esc7 had an MDK90 of 16 hrs, and Esc8 had an MDK90 of 19 hrs. These results showed that Esc7 and Esc8 had greater ethanol tolerance than SC5314 and that Esc6 had slightly higher tolerance over the first 12 hours of exposure.

### Screen of Deletion Strains Reveals Pathways Important for Ethanol Survival

Very little is known about factors that contribute to ethanol survival in *C. albicans* and therefore, a screen of transcription factor deletion mutants [17] was conducted. Strains were tested using the four-hour 10% ethanol exposure followed by plating on YPD-agar as described above. Strains were selected for analysis if the deleted transcription factor was homologous to an *S. cerevisiae* transcription factor with a role in ethanol survival, or if it regulated genes in pathways with a known or hypothesized role in ethanol survival.

Ethanol survival experiments were performed for 18 different homozygous transcription factor deletion strains and the wildtype strain, SN152 using six to twelve biological replicate cultures for each strain. Results are shown in Fig. 2. Some of the transcriptional regulators that are thought to be central to the *S. cerevisiae* ethanol response[18], e.g. *MSN4*, *MNL1 (scMSN2 analog*), and *CAP1* (*scYAP1 analog*) [18,19], were not necessary for ethanol survival in *C. albicans*. Other deletion strains had an altered ability to survive in 10% ethanol. One strain, deleted for *NDT80*, had a reduced ability to survive in ethanol, and four deletion strains (mutants lacking *TYE7, ACE2, YOX1,* or *CRZ1)* had an increased ability to survive in ethanol. *ACE2* and *CRZ1* knockout strains had particularly striking phenotypes with roughly 23 and 58-fold increases in ethanol survival respectively compared to WT (SN152).

**Figure 2:**
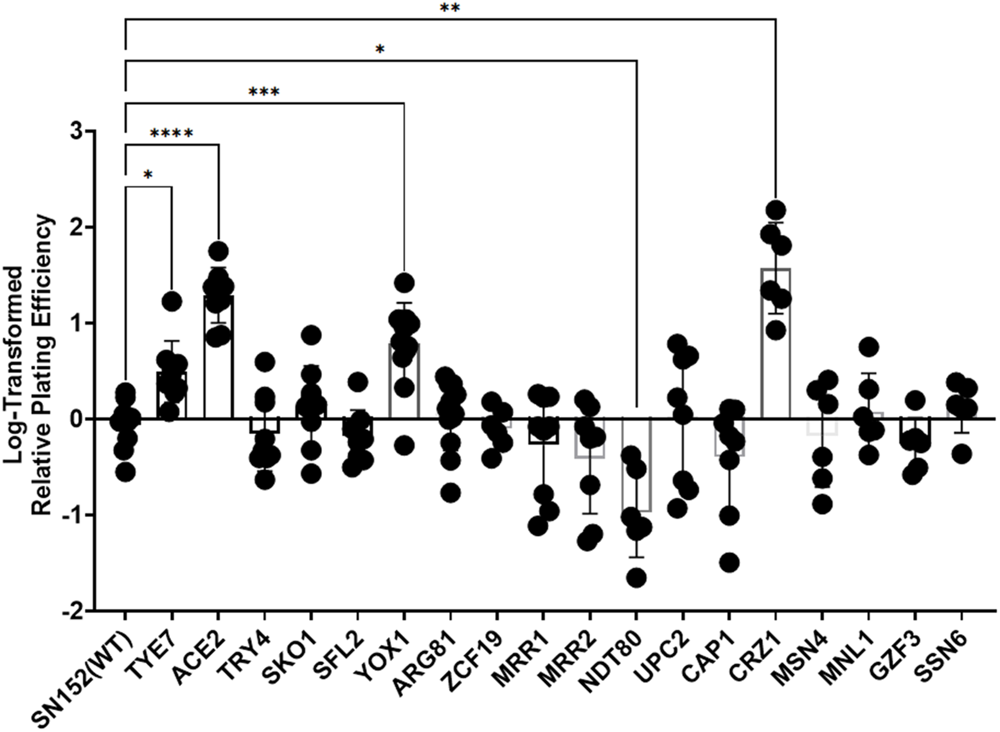
Screen of the Ethanol Survival of Transcription Factor Deletion Strains. Strains described in Homann, et al [17] were exposed to 10% ethanol for four hours as previously described. Relative survival was compared to SN152 (WT) as previously described. Relative survival was log-transformed and graphed. Eighteen homozygous deletion mutants are graphed with the name on the x-axis representing the deleted gene. Each point shows an individual biological replicate culture, bars show mean values, and error bars show standard deviation. A Brown-Forsythe and Welch ANOVA test was performed on log-transformed data to determine statistical significance; * = p < 0.0393; ** = p < 0.0016; *** = p < 0.0005; **** = p < 0.0001.

To identify critical pathways for ethanol survival, we used previous studies to ask whether pathways important for *S. cerevisiae* ethanol tolerance were regulated by these transcription factors. It was found that all 5 transcription factors have been implicated in regulation of the ergosterol pathway, either by co-expression studies, *ERG* promoter binding, or by computational prediction [19–23]. Additionally, four out of the five transcription factors have putative roles in biofilm formation, three in filamentous growth, two in cell-cell adhesion, two are involved with cell wall synthesis/organization, and two with susceptibility to azoles based on results described in the Candida Genome Database.

### *C. albicans* strains with elevated ergosterol levels exhibit increased ethanol tolerance

In *S. cerevisiae,* increased ergosterol content confers greater ethanol tolerance and in synthetic bilayers, increased ergosterol leads to lower membrane disruption [18,24–26]. Based on our findings from the transcription factor deletion mutant screen, we hypothesized that ergosterol is involved in *C. albicans* ethanol tolerance as well, and that Esc strains had alterations in ergosterol levels. We therefore analyzed the ergosterol content in the ethanol tolerant strains. SC5314 cells or Esc6, 7, 8 cells were cultured in medium A with 10% water or 10% ethanol for four hours and ergosterol was extracted and quantitated as previously described [27,28]. Ergosterol quantity was calculated relative to SC5314 in 10% H2O and is shown in Fig. 3A (8 biological replicates from four different experiments). From these data, a statistically significant increase in ergosterol content was seen in Esc7 in EtOH (p-value <0.0001, 2way ANOVA with Dunnett’s correction, mean = 1.77). Esc6 and Esc8 ergosterol content were not significantly different from SC5314. These data support the model that higher levels of ergosterol in Esc7 promoted higher ethanol tolerance.

**Figure 3:**
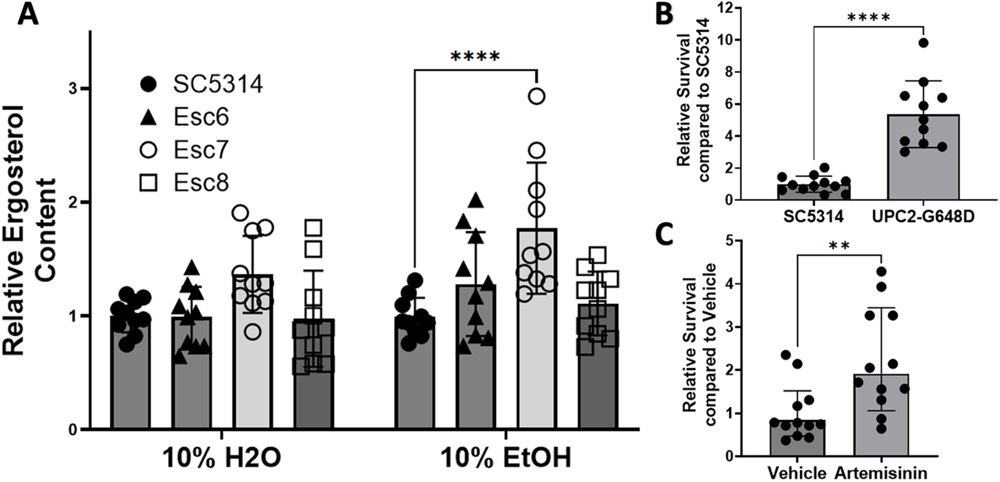
Cells with increased ergosterol content show increased ethanol tolerance. **A)** Cells were grown for 4 hours in medium A with 10% ethanol or water and harvested. Ergosterol was measured as a percentage ergosterol by mass normalized to wet-pellet weight calculated as previously described [27,29,30] and Materials and Methods). Relative ergosterol content was calculated by standardizing ergosterol % within each experiment to SC5314 in 10% H2O. 4 separate experiments with 10 total biological replicate cultures for each condition are shown. Symbols show individual replicate values relative to SC5314 within the condition and bars show means with standard deviation (****, p < 0.0001). Statistical significance calculated with 2way ANOVA with Dunnett’s multiple comparison, only significant comparisons are shown. **B)** Ethanol survival was measured after a 4-hr exposure of SC5314 or SC5314 with a *UPC2* gain of function mutation (G648D) [31]. Relative survival was standardized to SC5314 survival within each experiment. 11-12 biological replicates total from 3 different experiments are shown. One high outlier was removed from UPC2-G648D by ROUT (Q=1%). Symbols show individual replicate cultures and bars show mean with standard deviation (****, p < 0.0001). Statistical significance calculated by unpaired t-test. **C)** Ethanol survival from SC5314 cells pre-treated for 4 hrs with DMSO or DMSO with 100ug/mL of Artemisinin was measured. Each condition has 12 biological replicates. Ethanol exposure and standardization are the same as in B. Symbols show standardized ethanol survival of individual replicates and bars show median survival with error bars representing 95% confidence intervals. Statistical significance was tested using a Mann-Whitney test (**, p<0.0053).

We therefore tested whether elevating ergosterol content in *C. albicans* through other means would confer greater ethanol tolerance. One approach utilized an SC5314 strain with a homozygous *UPC2* gain of function mutation (G648D; kind gift of J. Morschhauser). *UPC2*_G648D_ results in increased ergosterol synthesis, and increased ergosterol content through upregulation of *ERG11* [31]. Four hour ethanol survival of the SC5314 WT strain [31] and homozygous *UPC2*_G648D_ mutant strain was determined as described previously. Relative survival compared to SC5314 was calculated. The homozygous *UPC2*_G648D_ mutant strain exhibited a five-fold increase in ethanol survival (Fig. 3B), showing that a mutation that increased ergosterol content [31] also increased ethanol tolerance.

For a second test of the hypothesis that increased ergosterol content increases ethanol tolerance, chemical stimulation of ergosterol synthesis with the anti-malarial drug artemisinin was performed. Incubation of *C. albicans* with 100ug/mL of artemisinin has been shown to increase ergosterol gene expression and lead to increased content of ergosterol in *C. albicans* cells [32]. Therefore, *C. albicans* cells were pretreated with 100ug/mL of artemisinin in DMSO or DMSO alone for four hours. Artemisinin or vehicle exposure was continued during the four-hour ethanol survival experiment. Survival of cells relative to SC5314 with vehicle was calculated (Fig. 3C). Stimulation of ergosterol synthesis with artemisinin led to a 2-fold increased ethanol survival. Therefore, cells with increased ergosterol content produced in several ways showed increased ethanol tolerance. Increased ethanol tolerance in Esc7 is likely due, at least in part, to increased ergosterol content in Esc7 cells.

### Altered gene expression in Esc strains

We next investigated gene expression differences between Esc7 and SC5314 during growth in medium A with ethanol versus medium A with water. SC5314 and Esc7 were cultured in 10% EtOH medium or 10% water medium for 4 hours and RT-qPCR was used to measure gene expression. We analyzed expression of genes of the ergosterol biosynthetic pathway that comprise roughly half of the biosynthetic pathway from lanosterol demethylation through ergosterol production [33] including *ERG11, ERG2, ERG4, ERG5, ERG6,* and *ERG25.* These genes were shown to have higher transcript abundance in Esc7 in comparison to SC5314 in 10% ethanol (Fig. 4A). Many of these genes were repressed in ethanol in SC5314 and less repressed in Esc7 in ethanol. These results were consistent with the increased concentrations of ergosterol in Esc7 compared to SC5314 since increased expression of *ERG11* was shown to increase ergosterol content in *C. albicans* [33].

**Figure 4:**
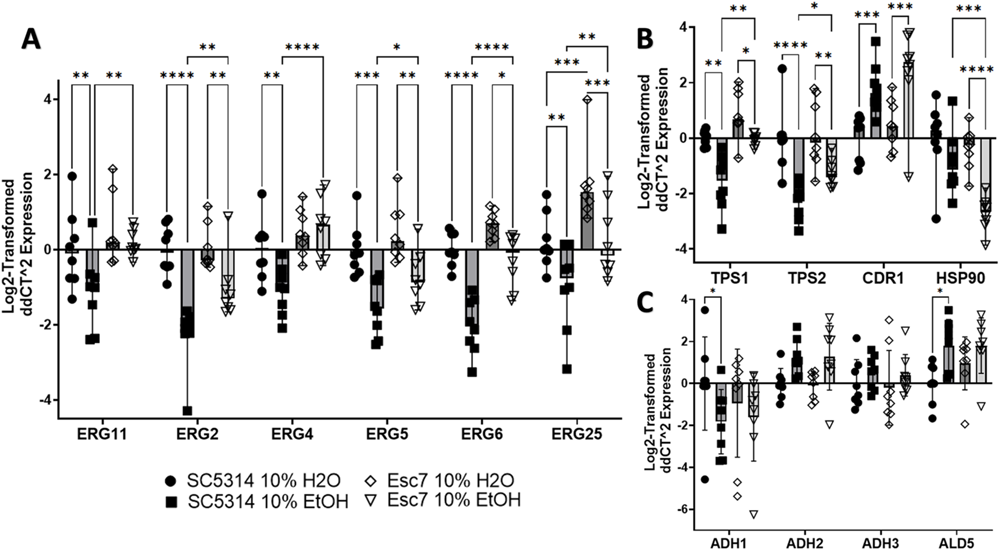
Differential gene expression in Esc7 versus SC5314 with and without ethanol. 8 biological replicate cultures were grown for 4 hours in medium A with 10% ethanol or water. Gene expression was measured by RT-qPCR. Expression was normalized to expression of *SPL1* (tRNA splicing gene that had unchanged expression in all conditions). Genes are indicated on the x-axis and Log_2_-fold change in expression is shown relative to SC5314 in 10% water on the y-axis. **A)** Genes of the ergosterol pathway. **B)** Genes/pathways implicated in *S. cerevisiae* ethanol tolerance. **C)** Genes in the ethanol catabolic pathway. Each symbol shows results from an individual culture, error bars show standard deviation, and statistical significance for each of the RT-qPCR figures was tested by Fisher’s LSD test. Only significant differences within one variable comparisons are shown. (*, p< 0.455, **, p< 0.0063, ***, p< 0.0004, ****, p< 0.0001)

Ethanol exposure has multiple effects that promote cell death including targeting the membrane, and producing protein misfolding, mitochondrial stress, and DNA damage [34–36]. In *S. cerevisiae,* increased expression of drug transporters and efflux pumps confers increased ethanol tolerance [37,38] and we confirmed upregulation of *CDR1* in SC5314 and Esc7 during exposure to ethanol as well as a trend towards greater expression in Esc7 compared to SC5314 (SC5314 vs Esc7 EtOH p-value: 0.2070, Fisher’s LSD test, mean value 1.51-fold higher) (Fig. 4B). Orthologs of genes involved in trehalose synthesis and ethanol tolerance in *S. cerevisiae, TPS1* and *TPS2* [18], were less repressed in Esc7 compared to SC5314 in 10% EtOH (Fig. 4B). Heat shock proteins (HSPs) and molecular chaperones are important for *S. cerevisiae* ethanol tolerance [39–41]. Interestingly, *HSP90* was significantly repressed in Esc7 in ethanol and was more repressed than in SC5314 (Fig. 4B). HSP90 is critical to many different stress responses, regulates Upc2 (a transcriptional regulator of ergosterol metabolism), and regulates antifungal resistance and tolerance [42–48]. Decreased expression of *HSP90* could lead to less programmed cell death during ethanol stress as has been observed with exposure to hydrogen peroxide [49].

Another pathway of interest is the ethanol catabolic pathway. This pathway has not been extensively studied in *C. albicans,* however, and some of the main players in this pathway were identified based on homology. Adh1 and Adh2 are thought to convert ethanol into acetaldehyde, Ald4 and Ald5 likely convert acetaldehyde into acetate, and Acs1 and Acs2 likely convert acetate into acetyl-CoA [50]. Several of the genes, e.g. *ADH2*, *ADH3*, *ADH4*, and *ALD5* were transcriptionally induced in ethanol in both SC5314 and Esc7 (Fig. 4C). Significant differences in expression of these genes between Esc7 and SC5314 were not observed by RT-qPCR.

A second approach to detect changes in gene expression that could contribute to ethanol tolerance, was through RNA-sequencing. Three biological replicates of SC5314 and Esc6, 7 and 8 were cultured in medium A with 10% ethanol or 10% water and harvested. RNA-seq was performed on RNA extracted from these samples as described in Materials and Methods and results are presented in **Supplemental Tables S1-S10**. Principal coordinate analysis showed that all Esc strains exhibited distinct transcriptional profiles in ethanol compared to SC5314 in ethanol (**Supplemental Fig. S4**). Interestingly, *ACE2* was the most reduced transcript in Esc7 compared to SC5314 in 10% ethanol and was 11.3 and 4.3 fold less abundant in Esc6 and Esc8 in ethanol compared to SC5314 in ethanol respectively, consistent with the elevated ethanol tolerance of the *ace2* deletion mutant (Fig. 2).

To determine what cellular pathways were altered in these strains, GO term analyses using the Candida Genome Database (CGD) were performed as previously described [51] to identify enriched categories of genes. Genes that were significantly up-regulated (log2-fold change >0.5 and adjusted p-value < 0.05) in Esc strains in ethanol compared to SC5314 in ethanol were analyzed (1700 genes in Esc6, 1334 genes in Esc7, and 1494 genes in Esc8). The top 20 processes based on corrected p-value identified by GO term process analysis are shown in **Supplemental Fig. S5**. Organonitrogen compound catabolic process was the only process that was shared in the top 20 for all strains.

For Esc7, processes involved with ergosterol biosynthesis (red arrows in Fig. 5) were enriched among the top 20 GO term processes, consistent with the analysis above. Esc6 and Esc8 showed other enriched pathways that could be involved in ethanol tolerance or response to ethanol exposure. Both Esc6 and Esc8 showed enrichment of genes involved with general catabolic processes. Both strains showed enrichment of processes pertaining to proteolysis and protein catabolism which could be involved with the denatured protein response caused by ethanol exposure. Esc6 also showed an enrichment of genes involved with autophagy, which has previously been implicated in the ethanol response of *S.* cerevisiae [52]. These findings suggest that changes to proteasomal degradation/proteolysis and autophagy could contribute to ethanol tolerance of *C. albicans*.

**Figure 5:**
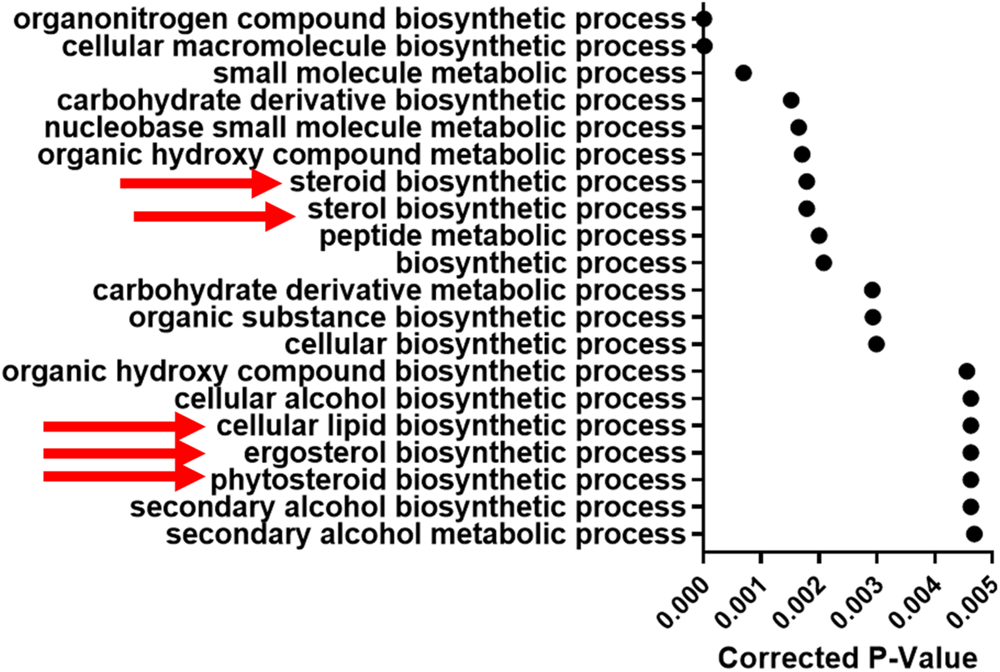
Differential gene expression in Esc7 versus SC5314 in 10% Ethanol. Transcripts expressed more highly in Esc7 in 10% ethanol compared to SC5314 were analyzed using GO term search on Candida Genome Database. Enriched processes are listed. The X axis shows corrected p-value for the top 20 hits (from most significant at top and descending statistical significance moving to the bottom) (triangles). Red arrows indicate processes related to lipid or ergosterol synthesis.

These findings highlight that many pathways are likely involved in ethanol tolerance in *C. albicans.* Some of the pathways involved in *S. cerevisiae* ethanol tolerance may not play a role in the strains described here (e.g. HSPs, ethanol catabolic pathway, etc.), although genes that do not change transcriptionally may also be regulated post-transcriptionally.

### Esc strains show chromosome copy number variation and genomic polymorphisms

Whole-genome sequencing of parental SC5314, Esc6, Esc7 and Esc8 was conducted to identify possible chromosome copy number variation or sequence polymorphisms in the ethanol tolerant strains. Aneuploidy is a well described mechanism of stress adaptation in *C. albicans* [53–55], as well as *S. cerevisiae* ethanol tolerance [56]. Genomic DNA from the strains was isolated and genome sequencing was conducted by Seq-Center (formerly MiGS) using paired-end Illumina next generation sequencing. Further details on sequencing and analysis workflow are described in Materials and Methods.

Changes in chromosome copy number were detected by uploading fastq files to the Yeast Mapping Analysis Pipeline (YMAP) [57]. YMAP analyzed average read depth across all chromosomes to give a read-depth based chromosome copy number estimation (Fig. 6). These images also give predictions for loss of heterozygosity (LoH) regions and show which regions are predicted to be composed of which alleles (A or B, AAB or ABB in trisomic regions) based on sequence homology to either of these alleles. Results show that Esc strains contain various aneuploidies, as well as certain LoH regions. Specifically, Esc6 has a trisomy of Chr1, Esc7 has a trisomy of Chr3, and a loss of the B allele of the short arm of chromosome R. Esc8 has a monosomy of Chr5 (only A allele) and trisomies of Chr6, 7, and 8. Growth in ethanol may select for aneuploid strains previously existing in the population or the presence of ethanol or byproducts of its catabolism may stimulate chromosomal instability.

**Figure 6:**
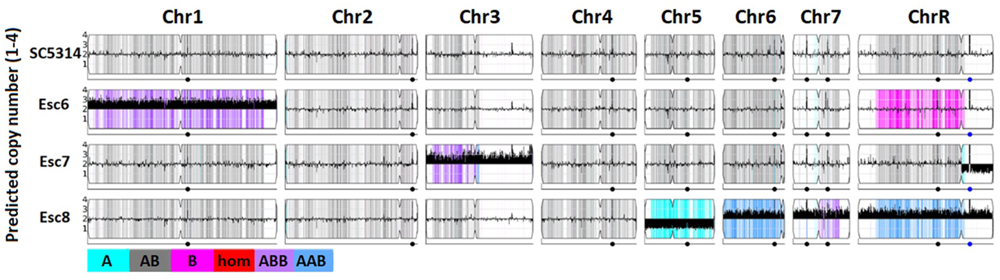
Chromosome copy numbers in Esc strains. Whole genome sequencing of SC5314, Esc6, 7, and 8 was conducted as described in Materials and Methods. Copy number estimates were determined based on read depth and estimated copy number is plotted along chromosomes using YMAP[57]. C. albicans is diploid so euploid chromosomes show an average copy number of 2 on the Y-axis as shown for SC5314. Thick black bars denote aneuploidies, shown by predicted copy number greater than or less than 2. Images from YMAP [57].

Additionally, we observed single nucleotide polymorphisms (SNPs) and insertions/deletions in the genome sequences of Esc6, Esc7, and Esc8 (**supplemental tables S11-S13**). There were 124 (in 64 different ORFs), 139 (in 83 different ORFs), and 135 (in 72 different ORFs) polymorphisms in protein coding regions in Esc6, Esc7, and Esc8 respectively. Polymorphisms were comprised mostly of putative heterozygous polymorphisms in all strains based on the allele depth ratio between the reference and alternate alleles. Heterozygous and homozygous polymorphisms were classified by creating a histogram of allele depth and calculating minima of a trimodal fit curve (minima = Esc6:0.1845,**0.8482**; Esc7:0.1753,**0.8716**; Esc8:0.1761,**0.8628**) (**Supplemental Fig. S6**). Further analysis focused on homozygous polymorphisms classified as variant calls with an allele depth ratio greater than the highest minimum value (bolded values above). Only one gene carried a homozygous polymorphism in all three Esc strains: *C6_02290C*, an uncharacterized gene containing a putative zinc binding domain. Other genes of interest with homozygous polymorphisms are *C4_04190C* and *C4_04200C* which are overlapping ORFs that both contain a putative acetyltransferase domain and *PDC2* which was polymorphic in two strains. *PDC2* encodes a homeodomain-containing transcriptional regulator which is involved in glucose metabolism [58]. The role of these genes in ethanol tolerance is unknown.

### Esc strains exhibit reduced susceptibility to fluconazole

Some of the chromosome aneuploidies observed in Esc7 and Esc8 have been previously observed in strains with altered susceptibility to the frontline antifungal fluconazole. Aneuploidies of chromosome 3, 5, 6, 7, and R have been implicated in azole resistance and tolerance [59–64]. Further, the ergosterol pathway, more specifically *ERG11,* is the target of azole class antifungals and increased expression of *ERG11* and *CDR1* confers fluconazole resistance [42,65–67].

We therefore examined the susceptibility of Esc strains to fluconazole and other antifungal drugs. Broth-microdilution assays were carried out using the standard CLSI protocol [68]. After 24-hour incubations, metabolic activity of the wells was measured with 2,3-bis(2-methoxy-4-nitro-5-sulfophenyl)-5-[(phenylamino)carbonyl]-2H-tetrazolium hydroxide (XTT) and MIC50 and MIC70 values were calculated as described in Materials and Methods (Fig. 7A, **Supplemental Fig. S7)**. Esc6 and Esc7 both exhibited fluconazole resistance phenotypes with elevated MIC50 values. Esc8 showed no alterations to MIC50 values but had increased MIC70 and trailing growth, suggesting altered fluconazole susceptibility. For amphotericin B and caspofungin, MICs were determined by visual determination of growth and all strains showed no changes in susceptibility to these antifungals (Fig. 7C).

**Figure 7:**
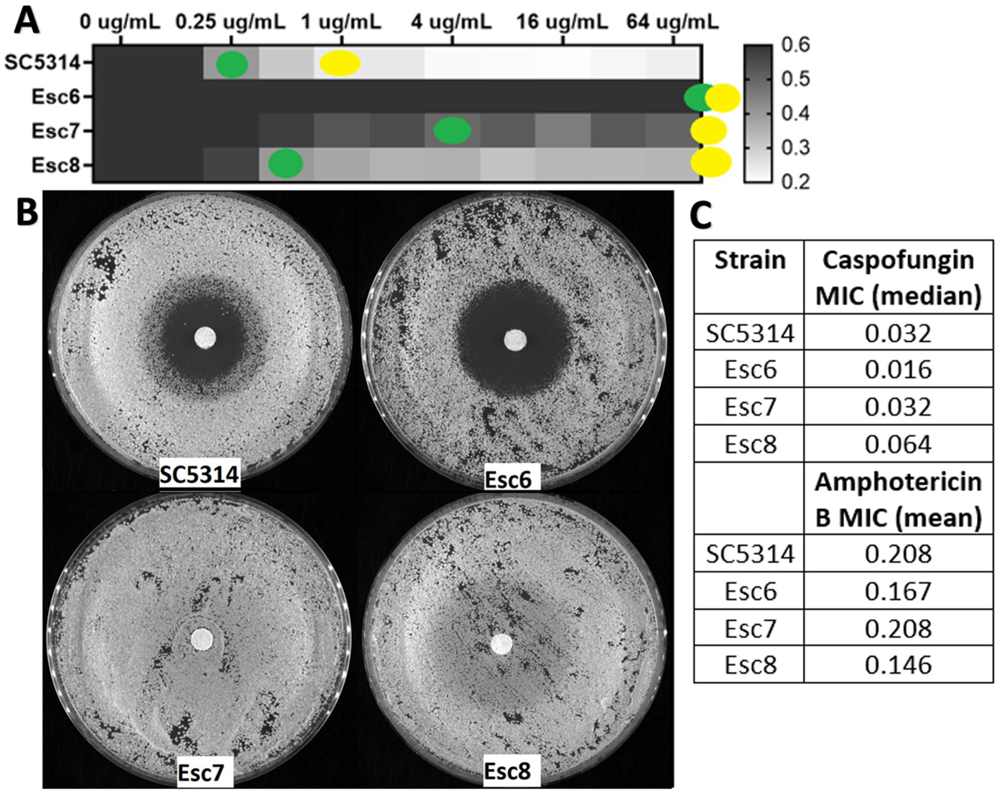
Esc strains exhibit altered fluconazole susceptibility. **A)** Strains were grown in liquid cultures containing increasing concentrations of fluconazole as described in Materials and Methods. Growth was detected after a 24 hour incubation using the XTT assay for metabolic activity. Metabolic activity relative to the 0ug/mL well is shown as a heat map. Yellow dots show the well that corresponds to MIC70 and green dots show MIC50. Median metabolic activity values from 6 biological replicates performed in 6 separate experiments are shown. **B)** Strains were grown to post exponential phase in YPD liquid and then used to seed the disk diffusion plates. Disk diffusion assays were performed on YPD agar with 10 cells/plate with 25ug fluconazole disks incubated for 48 hours at 30⁰C. Images were taken using a Syngene Imager. Representative images from an experiment repeated 4 times are shown. **C)** MIC values were determined using the same MIC protocol except MICs were determined by eye. The caspofungin MIC was determined from the mean of 4 biological replicate experiments for each strain and the amphotericin B MIC was determined from 3 biological replicate experiments. There were no significant differences between SC5314 and any of the Esc strains (testing for statistical significance performed on log-transformed data, Kruskal-Wallis test for caspofungin and One-Way ANOVA for amphotericin B with multiple comparisons for each)

Disk diffusion assays were performed following standard protocols with 25ug-loaded fluconazole filter disks [69,70]. Esc7 showed no zone of inhibition following a 48-hour incubation, while Esc8 showed a zone of inhibition with heavy growth within the zone (Fig. 7B). These phenotypes were similar for voriconazole, another azole antifungal drug (**Supplemental Fig. S8**). Esc7 showed no zone of voriconazole inhibition and Esc8 showed a zone of inhibition with heavy growth in the zone. These results are consistent with azole resistance and azole tolerance respectively. Interestingly, Esc6 showed a larger zone of fluconazole inhibition than SC5314 in the disk-diffusion assay but exhibited an elevated MIC for fluconazole in the broth microdilution assay. This type of discrepancy has been observed rarely and could suggest a non-canonical mechanism of reduced fluconazole susceptibility in this strain. In summary, all Esc strains tested exhibited altered susceptibility to fluconazole, showing that *C. albicans* can evolve fluconazole resistance and tolerance through ethanol exposure without previous exposure to antifungal drugs.

Due to this cross-tolerance/resistance, we compared the SNPs identified in Esc strains with SNPs previously identified in strains developing fluconazole resistance. Whole genome sequencing analysis of strains acquired by sequentially sampling patients during the acquisition of fluconazole resistance was reported by Ford, et al [69]. In this study, sequential isolates from patients with *C. albicans* oral candidiasis infections were obtained. In total 43 isolates from 11 patients were whole-genome sequenced. 7 of the 11 patients carried fluconazole resistant strains. Genome sequences and SNPs of these strains were analyzed throughout the sequential isolations and compared to SC5314 reference strains. From this study, 240 genes were observed with persistent polymorphisms, defined as polymorphisms observed in fluconazole resistant strains from at least 3 of the 7 patients, that were nonsynonymous, were not within a region of the genome with a loss of heterozygosity event and, after arising, were found in all subsequent isolates.

We determined how many of these 240 genes carried homozygous polymorphisms in Esc6, Esc7, and Esc8. There were 8 total homozygous SNP-carrying genes in Esc6 and 5 of these 8 polymorphic genes also carried persistent polymorphisms in strains analyzed by Ford et al. Of the 6 homozygous polymorphic genes in Esc7, 2 of these also carried persistent polymorphisms in the Ford, et al strains. Esc8 had 5 homozygous polymorphic genes of which 3 carried persistent polymorphisms in the Ford, et al strains. This overlap in genes was greater than expected by chance (p-values all < 0.0030; Chi-squared test). Genes that carry SNPs in Esc strains and Ford strains are shown in **Table 1**. All polymorphisms can be found in **supplemental tables S11-S13.** These results suggest an overlap in the selective pressures exerted by fluconazole and ethanol such that these two pressures select for similar mutational landscapes and aneuploidies. The similarities increase the likelihood that strains selected for higher ethanol tolerance will exhibit cross-tolerance/resistance to fluconazole.

**Table 1:** Polymorphic genes in Esc strains show overlap with genes associated with fluconazole resistance.

| Strain | Observed Number of Common Genes<sup>a</sup> | Expected Number of Common Genes<sup>b</sup> | Two-tailed P-value (Chi squared) |
| --- | --- | --- | --- |
| Esc6 | 5 | 0.3096 | <0.0001 |
| Esc7 | 2 | 0.2322 | 0.0029 |
| Esc8 | 3 | 0.1935 | <0.0001 |
| Homozygous Polymorphic Genes in Esc Strains |  |  |  |
| Esc6 | ALS3, EAP1, GZF3, RFG1, TBF1, C6_04190C, C7_01100C, CR_02930W |  |  |
| Esc7 | PDC2, C1_14630C, C1_07690C, C4_04190C, C4_04200C, C6_04190C |  |  |
| Esc8 | FMO2, PDC2, PGA55, C6_03520C, C6_04190C |  |  |
| Common Genes with Homozygous Polymorphisms in Esc strains and Persistent Polymorphisms in Ford, 2015 |  |  |  |
| Esc6 | C6_04190C, TBF1, CR_02930W, ALS3, EAP1 |  |  |
| Esc7 | C6_04190C, C1_07690C |  |  |
| Esc8 | C6_04190C, PGA55, FMO2 |  |  |

<sup>a</sup>240 previously identified open reading frames with persistent polymorphisms from Ford, et al, 2015 were compared with homozygous polymorphic open reading frames in Esc6, Esc7, and Esc8. The number of common open reading frames is listed for each strain.
<sup>b</sup>The frequencies of previously identified open reading frames with persistent polymorphisms from Ford, et al, 2015 and homozygous polymorphic open reading frames in Esc6, Esc7, and Esc8 were used to calculate the expected number of genes in common between these 2 gene sets.

## Discussion

In this study, we showed that repeated exposure of *C. albicans* cultures to ethanol results in selection for *C. albicans* small colony variants. The small colony variants that were studied exhibited increased ethanol tolerance as measured after a four-hour ethanol exposure and a longer term 24-hour ethanol exposure. Interestingly, Esc strains appeared to grow during the initial hours of the ethanol exposure and then lost viability at a similar rate to SC5314. This observation suggests that the evolved tolerance strategies in Esc strains were optimized for short ethanol exposures consistent with the four-hour exposures that were used during the selection. These mechanisms did not appear to provide much advantage during longer exposure times (i.e. 8-24 hours) as the rate of optical density decay was similar in SC5314 and all Esc strains in this time frame.

Little is known about how *C. albicans* responds to ethanol. A screen of homozygous transcription factor deletion strains supported the hypothesis that the ergosterol pathway is an important pathway that can impact ethanol survival in *C. albicans*. In general, ethanol kills cells through disruption of the lipid bilayer [71,72], protein denaturation [73], and intracellular membrane perturbations [74]. There is thinning of the bilayer through direct interdigitation of ethanol between membrane lipids, and ultimately denaturation of the membrane which will result in cell death [71,72]. We showed here that increased ergosterol content in *C. albicans* can confer increased ethanol survival. This is likely due to protection from the impacts of ethanol perturbation on the plasma membrane. Models of yeast plasma membranes showed that increased ergosterol content in the membrane reduced ethanol interdigitation and contributed to resistance against the membrane thinning effect of ethanol [72,75].

We asked whether *UPC2* was involved in the differential expression of *ERG* genes as Upc2 is a global regulator of ergosterol biosynthesis [76]. However, *UPC2* expression was decreased in Esc7 in 10% ethanol compared to SC5314 (**supplemental table S3**), and there were no SNPs in the *UPC2* orf or promoter sequence, suggesting that other novel transcriptional regulators of ergosterol synthesis are regulating *ERG* gene expression in Esc7. Additionally, post-transcriptional regulatory mechanisms could lead to greater activation of Upc2. For example in *S. cerevisiae* and *Candida* species, it has been suggested that HSP90 acts as a pedestal for Upc2 and binds it in an inactive form [77,78]. It is possible that similar mechanisms are at play in *C. albicans.* In Esc7, there is a higher ratio of *UPC2*/*HSP90* transcripts in Esc7 in 10% ethanol compared to SC5314 in 10% ethanol based on the RNA-Seq data (**Supplemental Fig. S9**). Regulation of the *UPC2/HSP90* ratio could lead to a greater proportion of Upc2 being in its active form, thus leading to greater ergosterol production. Furthermore, another transcriptional regulator of ergosterol gene expression, *ADR1* [79], also had no polymorphisms in any of the Esc strains.

Multiple mechanisms including lipid metabolism alterations, aneuploidy, increased production of stress responsive genes, and drug efflux pump expression have all been shown to confer greater ethanol tolerance in *S. cerevisiae* [18,56,80]. We hypothesized that *C. albicans* utilizes multiple mechanisms to increase ethanol tolerance and improve survival when exposed to relatively high concentrations of ethanol. Ethanol affects many different pathways in cells such as protein folding, lipid bilayer integrity, and mitochondrial and nuclear function [71–74,81]. In *C. albicans* specifically, ethanol also affects filamentation and biofilm formation [82].

Therefore, many different pathways are under selection during repeated exposure to ethanol and roughly 44% of all genes showed statistically significant alterations in gene expression in the 10% ethanol comparison of Esc7 and SC5314 (**supplemental table S3**). These large transcriptomic changes in Esc strains in ethanol indicate that many *C. albicans* pathways are likely impacted during ethanol exposure. The transcription factor deletion strains with altered ethanol tolerance affected many pathways and processes including biofilm formation, regulation of filamentation, cell-cell adhesion, and cell wall synthesis.

The Esc strains showed reduced susceptibility to the antifungal drug fluconazole and voriconazole but not to other antifungals. These observations suggest that at least part of the overlap of selective pressures between fluconazole and ethanol revolves around ergosterol. If a stressor impacts a pathway that an antifungal drug also targets, the stressor could theoretically select for drug resistant or tolerant strains. Even if there is no overlap in the phenotypic response to a stimulus and an antifungal, if the resistance mechanism involves aneuploidy, cross-resistance could arise due to increased dosage of genes. Previously this has been shown in *C. albicans* with pre-exposure to chemotherapeutic drugs leading to decreased caspofungin susceptibility [64]. We show similar but novel findings—*C. albicans* can evolve fluconazole resistance and higher tolerance after repeated exposure to a naturally occurring molecule, ethanol.

An unusual fluconazole resistance phenotype was observed for Esc6, defined as a susceptible disk-diffusion phenotype and a strong resistance phenotype observed by broth-microdilution. There have been well documented discrepancies between broth-microdilution assays and disk-diffusions [83]. However, discrepancies to the degree that we observed with Esc6 are uncommon. These observations suggest that Esc6 has developed a noncanonical mechanism of resistance, or that Esc6 shows an environment-dependent mechanism of resistance. Recently it has been shown that large discrepancies in susceptibility of vulvovaginal candidiasis (VVC) isolates to fluconazole can be observed in clinical testing laboratories depending on the pH of testing media. A pH of 4.5, which more accurately represents the pH of the vagina, is more likely to identify a VVC strain as resistant than pH 7. Testing at pH 4.5 more accurately explained treatment recalcitrance than susceptibilities obtained from testing at pH 7 [84]. Further studies of Esc6 will be needed to identify the basis for its fluconazole phenotypes.

Esc7 showed genetic alterations previously identified in fluconazole resistant strains including Chr3 trisomy [59–61] and higher expression of genes in the ergosterol pathway. These genetic changes have previously been implicated in fluconazole resistance [57–59]. However, Esc7 did not show large changes to *CDR1*, *CDR2*, or *MDR1* expression in the conditions we examined. Previously this was thought to be part of the mechanism of fluconazole resistance in strains with Chr3 trisomies as *CDR1, CDR2,* and *MRR1* reside on Chr3 [59,85].

Aneuploidies found in Esc8 included a monosomy of Chr5, and trisomies of Chr6, Chr7, and ChrR. ChrR trisomy has been found in fluconazole tolerant strains and shown to confer increased fluconazole tolerance [62–64]. Aneuploidies may contribute to the reduced fluconazole susceptibility of Esc8.

Aneuploidies, while likely part of the resistance and tolerance mechanisms of these strains, are not the only interesting genomic changes. Esc strains also carry SNPs in genes that were altered during the acquisition of fluconazole resistance in clinical isolates [69]. Likely, some of these SNPs confer a fitness advantage for Esc strains during growth in fluconazole. One of these polymorphic genes of interest is *C6_04190C* which encodes an uncharacterized orf with a putative zinc binding domain. *C6_04190C* was polymorphic in all Esc strains as well as the Ford, et al strains. Perhaps C6_04190C plays a role in zinc uptake or zinc sequestration, which could affect zinc-binding transcriptional regulators that are involved with fluconazole resistance such as Mrr1, Mrr2, Tac1, and Upc2 [86–88]. Modifying the pool of biologically available zinc could affect the activity of these various proteins and have effects on susceptibility to fluconazole.

In summary, we have shown that *C. albicans* can evolve increased ethanol tolerance following repeated exposure to ethanol. This is likely due to the involvement of many different pathways. Repeated ethanol exposure selects for Esc strains that have altered transcriptional responses to ethanol and altered genomic structure. We also showed that increased ergosterol content leads to enhanced ethanol survival. The mutational landscape of *C. albicans* Esc strains resembles the landscape of genes mutated in clinical isolates upon acquisition of fluconazole resistance. This finding highlights the overlap of selective pressures between the two stressors: fluconazole and ethanol. Consistent with this overlap, we demonstrated that strains repeatedly exposed to ethanol acquire fluconazole resistance and tolerance without previous exposure to antifungal drugs.

## Materials and Methods

### Strains and Culture Conditions

For the repeated ethanol exposure protocol, individual colonies of *C. albicans* strain SC5314 were inoculated into 10 mL of YPD and grown overnight at 30⁰C to post exponential phase (16-20 hours). 5 independent cultures were split into paired cultures to be exposed to media with ethanol or water. After overnight growth, cells were washed with PBS, counted on a hemocytometer and 10^6^ cells were inoculated from the overnight YPD cultures into 1 mL of medium A (Dulbecco’s Modified Eagle Medium (DMEM) with 10% fetal bovine serum and 1X non-essential amino acids) with ethanol or water in 2 mL screw cap microcentrifuge tubes. The concentrations of ethanol or water were 1% for the first exposure, 2% for the second exposure, 4% for the third exposure, 6% for the fourth exposure, 8% for the fifth exposure, and 10% for exposures six through eight (volume/volume). Cultures were incubated at 37⁰C in a 5% CO_2_ atmosphere for four hours to mimic physiological conditions. Following ethanol or water exposure, 10 uL of the medium A cultures were seeded into 10 mL YPD for recovery from the ethanol exposure and grown at 30⁰C until post-exponential phases were reached (1 day for passages in 6% ethanol or less or 2 days for passages in 8% and 10% ethanol). 100 uL of each population was plated on YPD agar and incubated for 2 days at 30⁰C following each medium A exposure to monitor population dynamics and normal sized and small sized colonies were counted by eye. Small colonies were defined as colonies estimated to be 50% or less in size compared to normal sized colonies.

Ethanol small colony (Esc) strains were isolated from different populations of cells at different time points in the repeated ethanol exposure protocol by picking colonies from the YPD agar plates. Populations 1-4 were populations repeatedly exposed to water and 5-8 were populations repeatedly exposed to ethanol. Esc6 was isolated from population 8 in the 6^th^ passage in ethanol (first exposure to 10% ethanol), Esc7 was isolated from population 6 in the 7^th^ passage in ethanol (second exposure to 10% ethanol), and Esc8 was isolated from the 5^th^ population in the 8^th^ passage in ethanol (third exposure to 10% ethanol). SC5314-UPC2-G648D was obtained from Dr. Joachim Morschhauser and has the following genotype: *UPC2-G648D::FRT/UPC2-G648D::FRT* [31].

For the ethanol screen with strains from the Homann collection [17], strains were obtained in a 96-well format from the Fungal Genetics Stock Center. Strains were then isolated from the 96 well plates and struck for isolation on YPD-agar plates. Individual colonies were isolated for each strain, and grown for 20 hours to post-exponential phase before use in the screen. The screen followed the same four-hour ethanol exposure protocol as described above.

### Ethanol Exposure to Test for Ethanol Survival

Strains were grown to post-exponential phase in YPD (16-20 hours) and then counted using a hemocytometer to seed 10^6^ cells in one mL of medium A with 10% H2O or 10% ethanol (vol/vol) using 95% ethanol in 2 mL screw cap tubes. Cultures were incubated for 4 hours in 37⁰C in a 5% CO_2_ atmosphere. Before incubation, 100 uL was sampled from the cultures and serially diluted onto YPD-agar plates for a starting CFU determination. Following 4 hours, cultures were sampled and diluted serially in 1X phosphate-buffered saline (PBS) and plated on YPD agar for a final CFU determination for the water and ethanol cultures. YPD-agar plates were incubated for 2 days at 30⁰C and counted. The final CFU from the ethanol cultures was divided by the final CFU from the water cultures to determine percentage survival and then calculated as a survival relative to SC5314 within each separate experiment. For the screen of transcription factor deletion mutants, these strains were compared to their parent strain, SN152, for a determination of relative survival.

For the extended incubation, the same conditions were used as above except cultures were incubated in 37⁰C incubators with ambient CO_2_ concentrations. Two separate cultures for each biological replicate were set up for each individual experiment. The first set of cultures was sampled at 0, 2, 4, and 6 hours and the second set was sampled at 8, 10, 12, and 24 hours. Survival was calculated relative to starting CFU obtained at timepoint 0 hours.

### RNA isolation, RT-qPCR, and RNA-sequencing

Cultures of SC5314 or Esc strains were grown to post-exponential phase and 10^8^ cells for each were seeded into 1 mL cultures of medium A with 10% H_2_O or EtOH and incubated for 4 hours at 37⁰C with 5% CO_2_. Immediately following the incubation, cells were centrifuged at 6000 rpm for 1 min and supernatant was removed. Cells were then resuspended in 1 mL of RNA-later (Fisher cat. AM7021) and immediately frozen at −80°C. Cells were thawed on ice and lysed as previously described by bead-beating [89]. RNA was isolated using the Invitrogen Purelink RNA mini kit (Thermo Fisher cat: 12183-025) with the on-column DNase treatment (Thermo Fisher cat: 12185-010). RNA was eluted in 40uL of ultrapure H_2_O and RNA yield and purity were assessed on a Nanodrop.

For RT-qPCR, cDNA was synthesized using SSIII reverse transcriptase (Thermo cat: 18080-085) following manufacturer’s protocol with approximately 1 ug starting RNA concentrations. Oligo-dT primers were used for each cDNA synthesis reaction. Eight biological replicates for each strain and condition were used. cDNA was diluted 1:5 in ultrapure H_2_O and 1 uL was used per reaction for RT-qPCR. SYBR green master mix (Fisher cat: 4334973) was used and qPCR reactions were run on a Roche Lightcycler 480 II using standard two-step reaction procedures. Controls without cDNA were run for each primer and all primer pairs either did not yield product, or yielded Cp values that were at least 10 cycles greater than the sample Cp value. Melting curves were analyzed to ensure only one peak was seen and primer dimerization was not observed in any of the curves analyzed. Cp values were normalized using *SPL1*, encoding a tRNA splicing enzyme that showed no significant changes in gene expression in all conditions. The fold change in gene expression relative to SC5314 in 10% H2O was then calculated and graphed by 2^-ΔΔCT^ values [90]. Primers are shown in **supplemental table S14**. Primer sequences were either used from previously published work or were designed using NCBI primer blast. Primer design used the A allele sequence of the target gene, *C. albicans* strain SC5314 reference genome, PCR product size of 90-250bp, optimized primer length of 28 nucleotides, primer melting temperatures of 53-63⁰C with 56⁰C optimized temperature and a max primer pair difference of 5⁰C. Purified PCR products were sequenced to verify target specificity.

For RNA-seq, three biological replicates of SC5314, Esc6, Esc7, or Esc8 following exposure to medium A with 10% water or 10% EtOH were sent for sequencing. All samples sequenced had RNA integrity scores of 9.2 or greater. cDNA libraries were made by Genewiz and RNA-seq was performed by Genewiz as previously described [70] with the following alterations: gene expression was considered significant only if adjusted-P-value (calculated by Benjamini-Hochberg test) was less than 0.05 and log_2_-fold change was greater than the absolute value of 0.5. RNA-sequencing was aligned to *Candida albicans* SC5314 (assembly ASM18296v3) as the reference genome. Gene ontology of significant hits were assessed using Candida Genome Database Go term finder as has been previously described [51] (http://candidagenome.org). For gene expression differences, the three biological replicates from each condition were averaged and significant differences were compared by relative transcript abundance differences.

### Ergosterol Quantitation

Total cellular ergosterol was quantitated as described previously [27,29,30]. 10^8^ cells of SC5314 or Esc strains were incubated for 4 hours at 37⁰C with 5% CO_2_ in medium A with 10% ethanol or 10% H_2_O. After the four-hour incubation, cells were pelleted and washed twice in sterile PBS. Following washing, supernatant was removed, and the wet pellet weight of the cells was measured. 2 mL of alcoholic 25% potassium hydroxide was added to each tube for saponification and pellets were resuspended by vortexing for 1 min, transferred to borosilicate glass tubes, and incubated in an 85⁰C water bath for 1 hour. Following the incubation, tubes were allowed to cool for 10 mins at room temperature and 2 mL of spectroscopy grade n-heptane (Sigma 1043660500) and 667 uL of sterile deionized water were added to each tube and vortexed for 3 mins. The organic fraction was removed from the mixture and diluted 1:2 in 100% ethanol (Sigma cat: E7023-1L). Scanning absorbance values were then taken between 230nm and 290nm on a Nanodrop spectrophotometer. Calculation of the amount of ergosterol and amount of dehydro-ergosterol (DHE) normalized to wet pellet weight was performed as described previously [43,82,83]. In summary, the calculation was **equation 1**: % (ergosterol + DHE) = [(A_282_/290) x dilution factor] / pellet weight. **Equation 2**: % DHE = [(A_230_/518) x dilution factor]/pellet weight. % ergosterol = **Equation 1 – Equation 2**. The dilution factor is the dilution factor used for diluting the sterol-containing heptane solution with ethanol, 290 = E for ergosterol at 282 nm, and 518 = E for DHE at 230 nm [27].

### Whole-Genome Sequencing

Genomic DNA from SC5314 and Esc strains was extracted and sent to SeqCenter for next-generation sequencing. SC5314 and Esc6 yielded consistent colony sizes so these strains were struck out on YPD-agar and isolated colonies were picked, grown to post-exponential phase and cells were pelleted. Esc7 and Esc8 have more unstable small colony phenotypes, therefore, cells were plated on a YPD-agar plate grown for 3 days at 30⁰C and small colonies were picked and added to 100uL of sterile PBS. This was then used to seed 4 new plates and performed three separate times. The plates with the lowest reversion rates were used for genome preps by collecting small colonies from these plates and adding them to 5 mL of sterile PBS. These tubes were then vortexed, and cells were pelleted. Genomes were extracted using bead-beating in phenol chloroform followed by chloroform extraction and ethanol precipitation as described previously [91]. DNA was purified further by the Qiagen DNeasy kit using manufacturer’s protocol.

Whole-genome sequencing was performed by SeqCenter by paired end NGS Illumina sequencing of 151 bp reads. Quality control and adaptor trimming were performed with bcl-convert [92], and reads were mapped to SC5314 reference genome using bwa [93]. Genome coverage from the sequencing runs ranged from 66X to 83X based on total number of reads across the SC5314 reference genome length. GATK’s Markduplicates was used to remove duplicate reads in the alignment and GATK’s Haplotype caller was used to call variants in the alignment [94]. Variants with a QD < 2, or MQ < 40, or MGRankSum < -12.5, or ReadPosRankSum < -8, or FS > 60.0, or SOR > 3 were filtered out. Sequence variants were compiled into an excel document by SeqCenter. Due to the large number of sequence variants obtained, only homozygous sequence variants of Esc strains compared to the parental SC5314 strain sequence were analyzed in depth. Homozygous sequence variants were determined by constructing a histogram of allele depth and calculating minima of a trimodal fit curve (**Supplemental Fig. S6**). Homozygous polymorphisms were classified as variant calls with an allele depth ratio greater than the high minima value. Heterozygous polymorphisms were classified as those with allele depth ratios between the two minima values. Polymorphisms with allele depth ratio lower than the high minima and less than a total read depth of 10 were excluded from analysis. For the remaining putative homozygous polymorphisms, pre-existing heterozygous polymorphisms in the B allele were excluded by examining the location of the polymorphism in a multiple sequence alignment of the A and B alleles of the gene.

Chromosome copy number variations were analyzed using the Yeast Mapping Analysis Pipeline [57]. Fastq files were uploaded to YMAP and YMAP calculated the average read depth across the chromosomes mapping to SC5314 as the reference genome. YMAP thus generates chromosome copy number predictions based on the read depth across the genome. These predictions were graphed as predicted copy number across chromosomes of the individual strains relative to a ploidy of 2. Changes in ploidy were shown by thick black bars deviating from the starting ploidy line at 2.

### Disk-Diffusion and Broth-Microdilution Assays

For the disk-diffusion assay, cells were grown to post-exponential phase in liquid YPD medium and 10^5^ cells per plate were spread onto YPD-agar. Then 6mm diameter Whatman #1 filter disks impregnated with 25ug of fluconazole or 1ug of voriconazole were placed in the center of the YPD agar. These plates were incubated for 48 hours at 30⁰C to assess the zone of inhibition and the fraction of growth within the zone. Images of plates following incubation were taken on a Syngene gel-doc imager using white light, black background, and identical iris, zoom, and focus settings for each plate. Representative images of each strain are shown. Eight biological replicates of each strain were tested in four different experiments for fluconazole or 6 biological replicates in three different experiments for voriconazole.

For the broth-microdilution assay, cells were grown as stated above and 10^4^ cells/well were used for each strain and experiment. Standard CLSI protocol was used for this experiment. Cells were diluted in RPMI, 165 mM MOPS, 2% glucose, pH 7.0. Antifungals were serially diluted 2-fold—yielding final concentrations from 64 ug/mL down to 0.125 ug/mL for fluconazole, 2 to 0.015ug/mL for caspofungin or 4 to 0.03ug/mL for amphotericin B. In each row, a no-cell well was used as a blank and sterility control, and a no-drug control was used to calculate 100% metabolic activity. Plates were incubated for 24 hours at 35⁰C. Growth was determined by visual growth for amphotericin B and caspofungin or by measurement of metabolic activity for fluconazole. For the latter measurement, medium was carefully removed from each well and 100 uL of 1 mg/mL of 2,3-bis(2-methoxy-4-nitro-5-sulfophenyl)-5-[(phenylamino)carbonyl]-2H-tetrazolium hydroxide (XTT) (Alfa-Aesar) in PBS with 4 uL of menadione per/100 uL was added to each well. The plates were incubated for one hour at room temperature and then 50 uL of solution from each well was removed and read at 492 nm to determine metabolic activity of each well. The no-drug well averaged between the two duplicates was used to calculate the OD492 that indicates a 50% reduction. The concentration of drug that yielded a reduction in metabolic activity immediately less than 50% of the no-drug well was considered the MIC50. The heat map of the metabolic activity of these wells is shown as the medians of 6 biological replicates for each strain in Fig. 7 or shown as individual values in **Supplemental Fig. S7**. Heat map was made using GraphPad Prism.

## Author Contributions

Conceptualization, A.W.D. and C.A.K.; investigation, A.W.D.; writing—original draft preparation, A.W.D.; writing—review and editing, C.A.K., A.W.D.; funding acquisition, C.A.K. All authors have read and agreed to the published version of the manuscript.

## Acknowledgements

We thank Dr. Joachim Morschhäuser (Universität Wurzburg) for the kind gift of the *UPC2*_G648D_ mutant strain, Dr. Albert Tai for help with the whole-genome sequencing analysis, and Drs. Jesus Romo, Paola Zucchi, and Ashlee Junier for helpful discussion. We would also like to thank Jeyra Perez Lozada and Dayana Alay Gonzalez for technical assistance.

## Data Availability Statement

Whole-genome sequencing fastq files are available on the NCBI sequence read archive (project: PRJNA1018434). Gene expression matrices are available on the Genome Expression Omnibus (GEO project: GSE243757).

## Supporting Information

**Supplemental Figure S1: Quantification of small colony variants throughout passaging of four separate populations of cells that were exposed repeatedly to increasing concentrations of ethanol in medium A or repeatedly exposed to water in medium A.** Quantification of small colony variants throughout passage numbers of four separate populations of cells that were exposed repeatedly to increasing concentrations of ethanol in media A or repeatedly exposed to water in media A. Green squares were repeatedly exposed to ethanol and black dots were repeatedly exposed to water. Y-axis shows % small colonies relative to total number of colonies. Bars represent the mean values of the populations at that passage number. Small colonies were classified as less than 50% of the size of the normal sized colonies on a plate by eye.

**Supplemental Figure S2: Enc and Wnc survival in 10% ethanol**. Enc and Wnc survival in 10% ethanol. Colonies exposed for 4 hours in 10% ethanol or 10% water and plated on YPD-agar to count final colonies. The percentage of final colonies in 10% ethanol compared to 10% water were calculated and used to calculate a relative survival compared to SC5314 within each experiment (y-axis). Normal-sized colonies from ethanol populations (Enc) or water population (Wnc) were used. One-way ANOVA with Welch’s correction was used for statistical testing with multiple comparisons. Median values with 95% confidence interval are plotted for 8-16 replicate cultures from 2-4 separate experiments for each strain.

**Supplemental Figure S3: Growth curves and doubling times of Esc strains and normal-sized colony strains in YPD.** Isolated colonies were picked and grown for 20 hours to post-exponential phase and seeded into YPD in 96 well plates at an approximate OD600 of 0.05. Cells were grown for 48 hours at 30⁰C with constant shaking in a Biotek Epoch2 plate reader with OD600 read every 30 minutes. Three biological replicate cultures from each strain were used to graph the growth curves. Growth curves show the mean and standard deviation of each timepoint. The wildtype, background strain: SC5314, a culture from the repeat media-A (with no ethanol exposure) is shown as RepDMEM-2, and a normal sized colony from a repeat ethanol exposure population is shown as RepEtOH-8. Esc6, Esc7, and Esc8 are all shown as well.

Timepoints 0-24 hours were used to calculate doubling times by performing a nonlinear regression in GraphPad Prism with the exponential plateau fit to find the rate constant: k. All R^2^ values were above 0.9331. k values were then used to calculate doubling times with the formula LN(2)/k. These were standardized to the SC5314 doubling times and then plotted in this figure as standardized doubling times.

**Supplemental Figure S4: Principal component analyses for SC5314 vs Esc6, Esc7, or Esc8 in 10% ethanol from RNA sequencing**. Principal component analyses for SC5314 vs Esc6, Esc7, or Esc8 in 10% ethanol each from RNA sequencing. PC1 variance ranges from 81%-93% and PC2 variance ranges from 4%-9%. SC5314 is shown in red dots in each and Esc strains are shown in blue. Each dot represents a separate biological replicate culture that was sequenced.

**Supplemental Figure S5: Gene Ontology Terms of the Top 20 Processes of Significantly Enriched Transcripts in Esc strains exposed to ethanol compared to SC5314 exposed to ethanol**. Gene Ontology Terms of the Top 20 Processes of Significantly Enriched Transcripts in Esc strains exposed to ethanol compared to SC5314 exposed to ethanol. Transcripts that were significantly enriched in Esc strains were used to determine GO term process clustering on Candida Genome Database GO term finder. The top 20 hits from each strain were added to the graph and shown on y-axis. Esc6 bars are shown in blue, Esc7 bars are shown in black, and Esc8 bars are shown in orange. X-axis shows the cluster frequency of each of these processes relative to the total number of genes in the respective sets of enriched transcripts used for the analysis.

**Supplemental Figure S6: Histogram Fit Curve of Esc6 Allele Depth Ratio.** Histogram fit curve of Esc6 allele depth ratio is shown. Esc6 is used as an example of what was done for all of the strains that were sequenced. Binning was performed by calculating the number of genes that had allele depth ratios within bins of 0.05. The bin of 0-0.05 was set to an arbitrary number as this represents homozygous for the reference strain and should not be called in the sequence variant report. Then a trimodal fit was performed in excel and minima values were calculated and used for further determination of heterozygous and homozygous polymorphisms.

**Supplemental Figure S7: MIC50 values from broth-microdilution experiments of SC5314 and Esc6, Esc7, and Esc8**. MIC50 values from broth-microdilution experiments of SC5314 and Esc6, Esc7, and Esc8. Six separate biological replicate experiments were conducted and plotted. MIC50 values were determined for each by measuring metabolic activity (XTT) and determining the well that represented a 50% reduction in metabolic activity. Geometric means with geometric SD are graphed and ordinary One-way ANOVA was performed for statistical significance.

**Supplemental Figure S8: Voriconazole Disk-Diffusion Images of SC5314 and Esc Strains.** SC5314 and Esc strains (-1 and -2 indicate biological replicate cultures) were grown to post exponential phase and then seeded onto YPD agar at a concentration of 10^5^ cells per plate. A 1 ug disk of voriconazole was placed in the center of the plate. These plates were incubated at 30⁰C for 48 hours and then imaged with the same settings and converted to black and white images.

**Supplemental Figure S9: UPC2 to HSP90 Ratio of SC5314 vs Esc7 in different conditions**. UPC2 to HSP90 Ratio of SC5314 vs Esc7 in different conditions. Arbitrary transcript values from RNA-seq data were used to calculate transcript ratios of UPC2 to HSP90 and were plotted (y-axis). Ordinary one-way ANOVA used for statistical significance. NS = not significant. * = p-value of 0.0377.

Supplemental Table S1 Transcripts showing significant changes in abundance between SC5314 with ethanol vs water medium by RNASeq

Supplemental Table S2 Transcripts showing significant changes in abundance between Esc7 with ethanol vs water medium by RNASeq

Supplemental Table S3 Transcripts showing significant changes in abundance between Esc7 with ethanol vs SC5314 with ethanol medium by RNASeq

Supplemental Table S4 Transcripts showing significant changes in abundance between Esc7 with water vs SC5314 with water medium by RNASeq

Supplemental Table S5 Transcripts showing significant changes in abundance between Esc6 with ethanol vs water medium by RNASeq

Supplemental Table S6 Transcripts showing significant changes in abundance between Esc8 with ethanol vs water medium by RNASeq

Supplemental Table S7 Transcripts showing significant changes in abundance between Esc6 with ethanol vs SC5314 with ethanol medium by RNASeq

Supplemental Table S8 Transcripts showing significant changes in abundance between Esc8 with ethanol vs SC5314 with ethanol medium by RNASeq

Supplemental Table S9 Transcripts showing significant changes in abundance between Esc6 with water vs SC5314 with water medium by RNASeq

Supplemental Table S10 Transcripts showing significant changes in abundance between Esc8 with water vs SC5314 with water medium by RNASeq

Supplemental Table S11 Esc6 Genomic Polymorphisms Supplemental Table S12 Esc7 Genomic Polymorphisms Supplemental Table S13 Esc8 Genomic Polymorphisms Supplemental Table S14: Primers used in this study.

