## Supplementary figures and images for "Selection of Ethanol Tolerant Strains of *Candida albicans* by Repeated Ethanol Exposure Results in Strains with Reduced Susceptibility to Fluconazole"

### Supplemental Figure S1

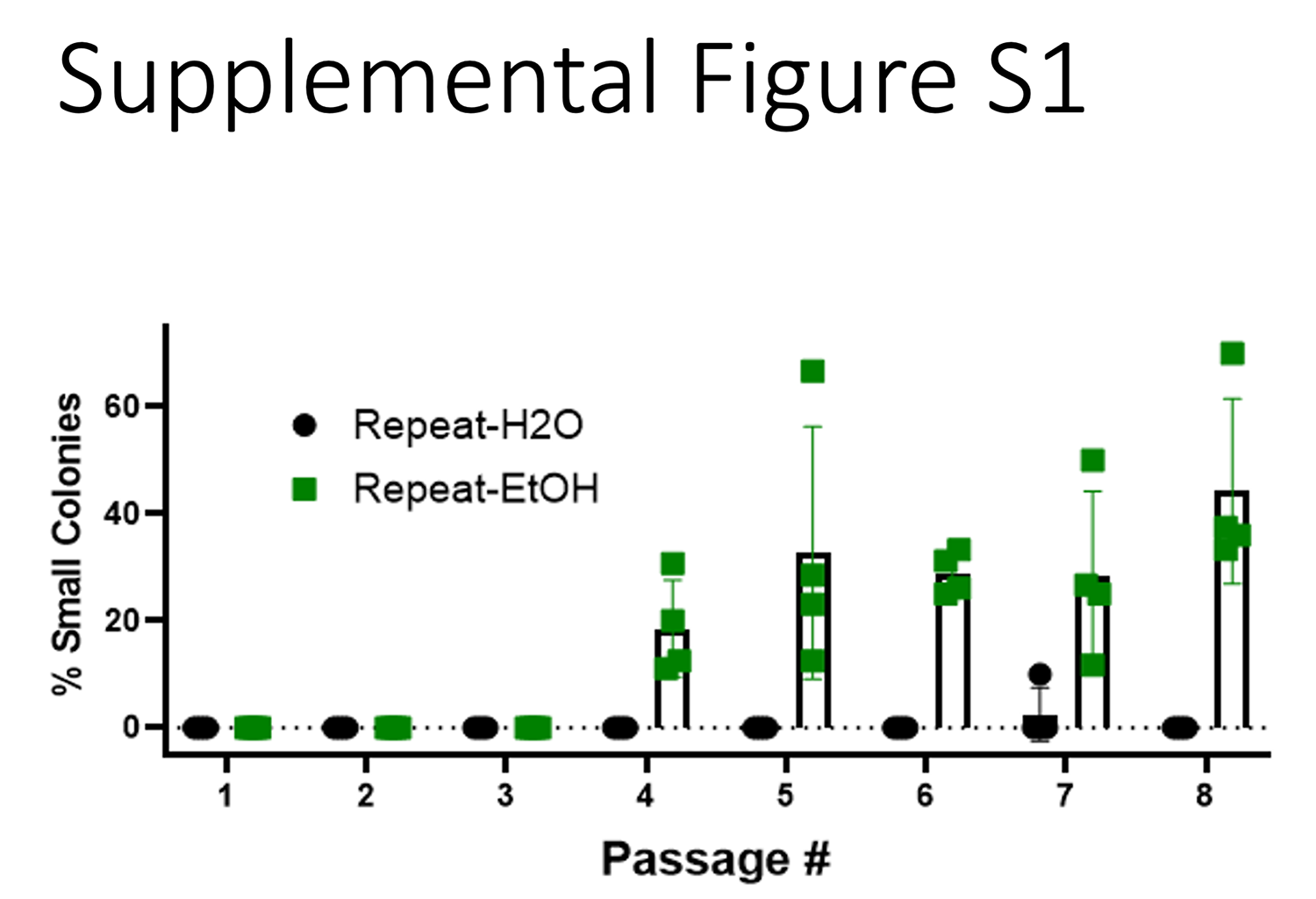

### Supplemental Figure S2

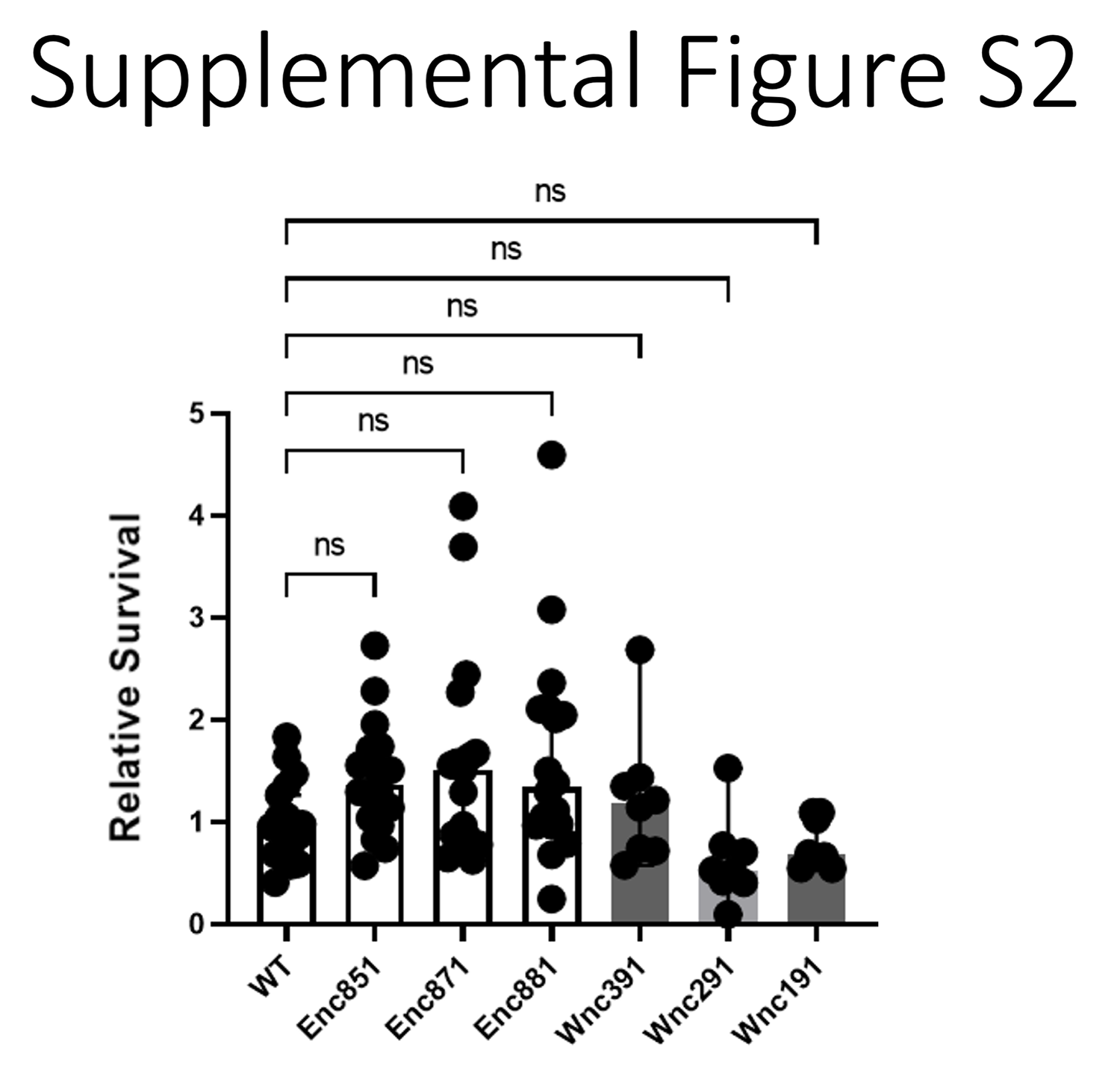

### Supplemental Figure S3

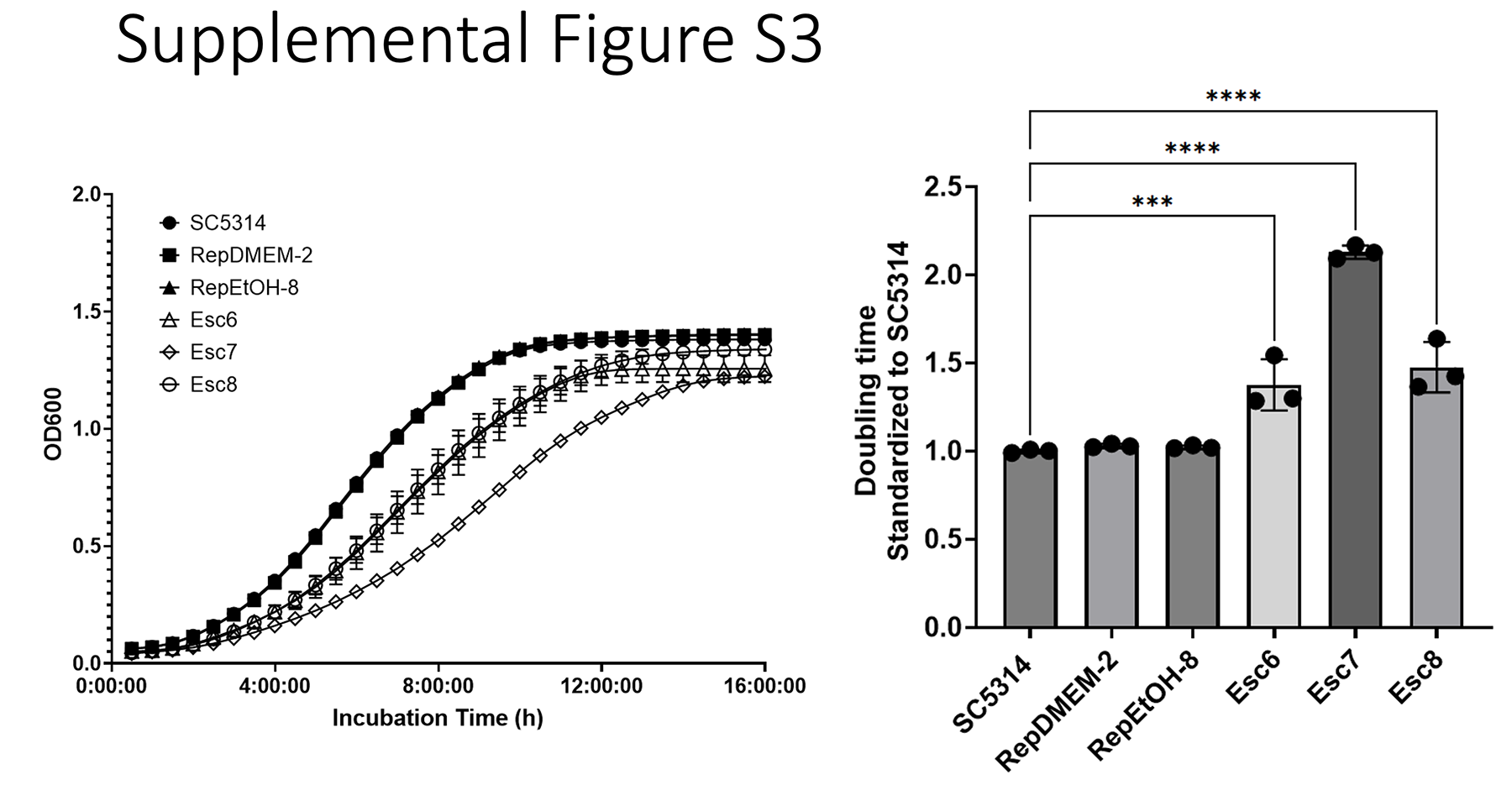

### Supplemental Figure S4

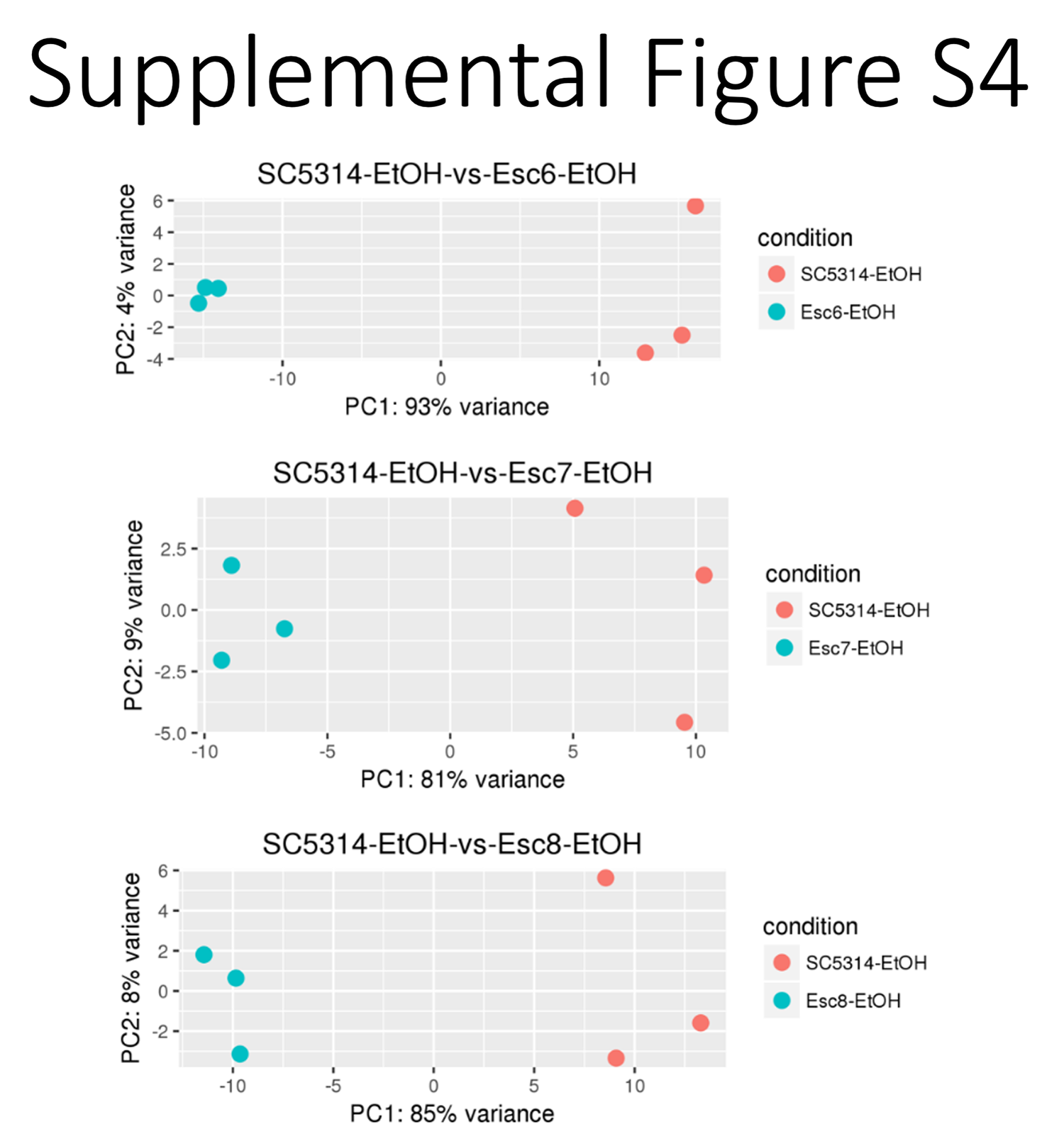

### Supplemental Figure S5

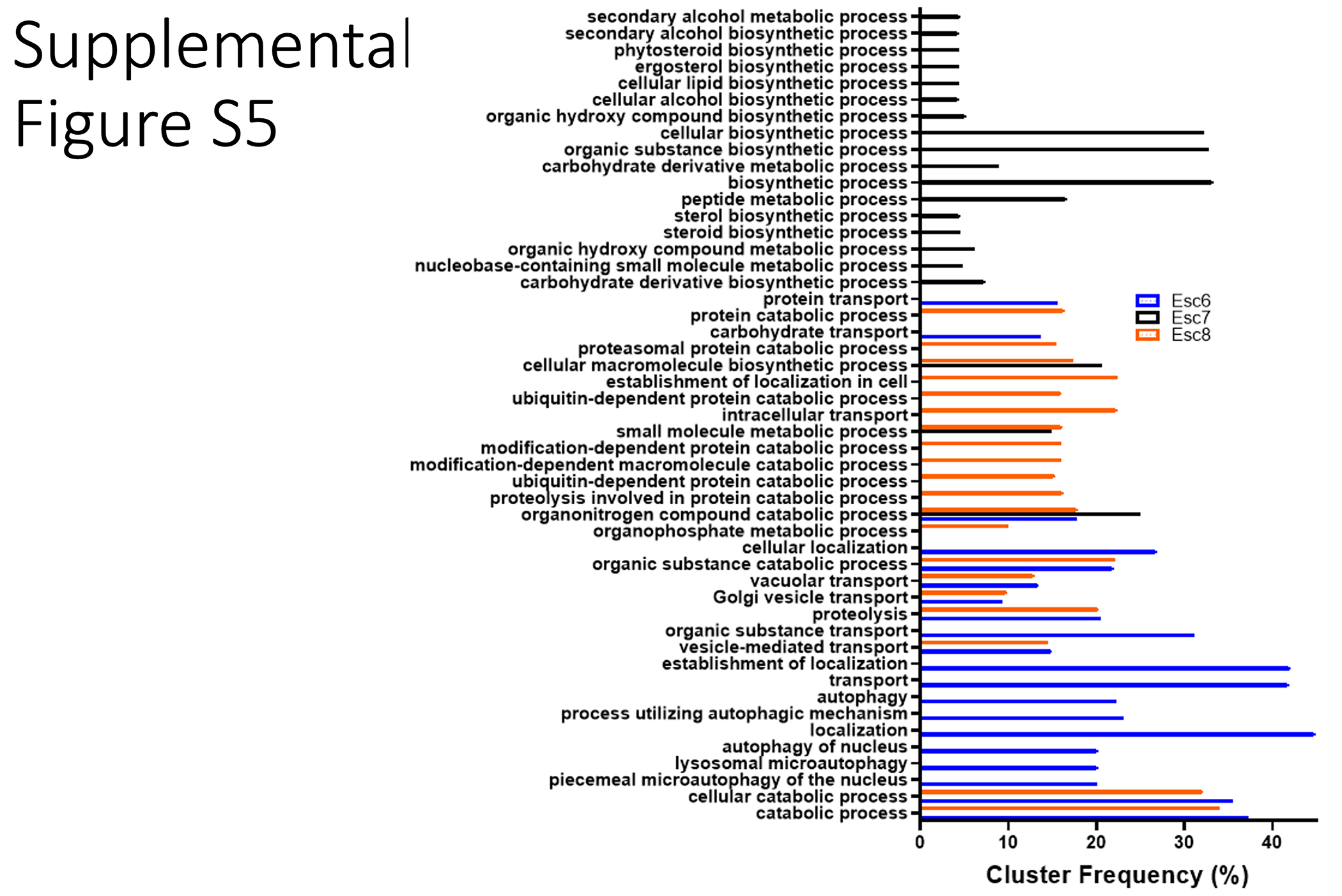

### Supplemental Figure S6

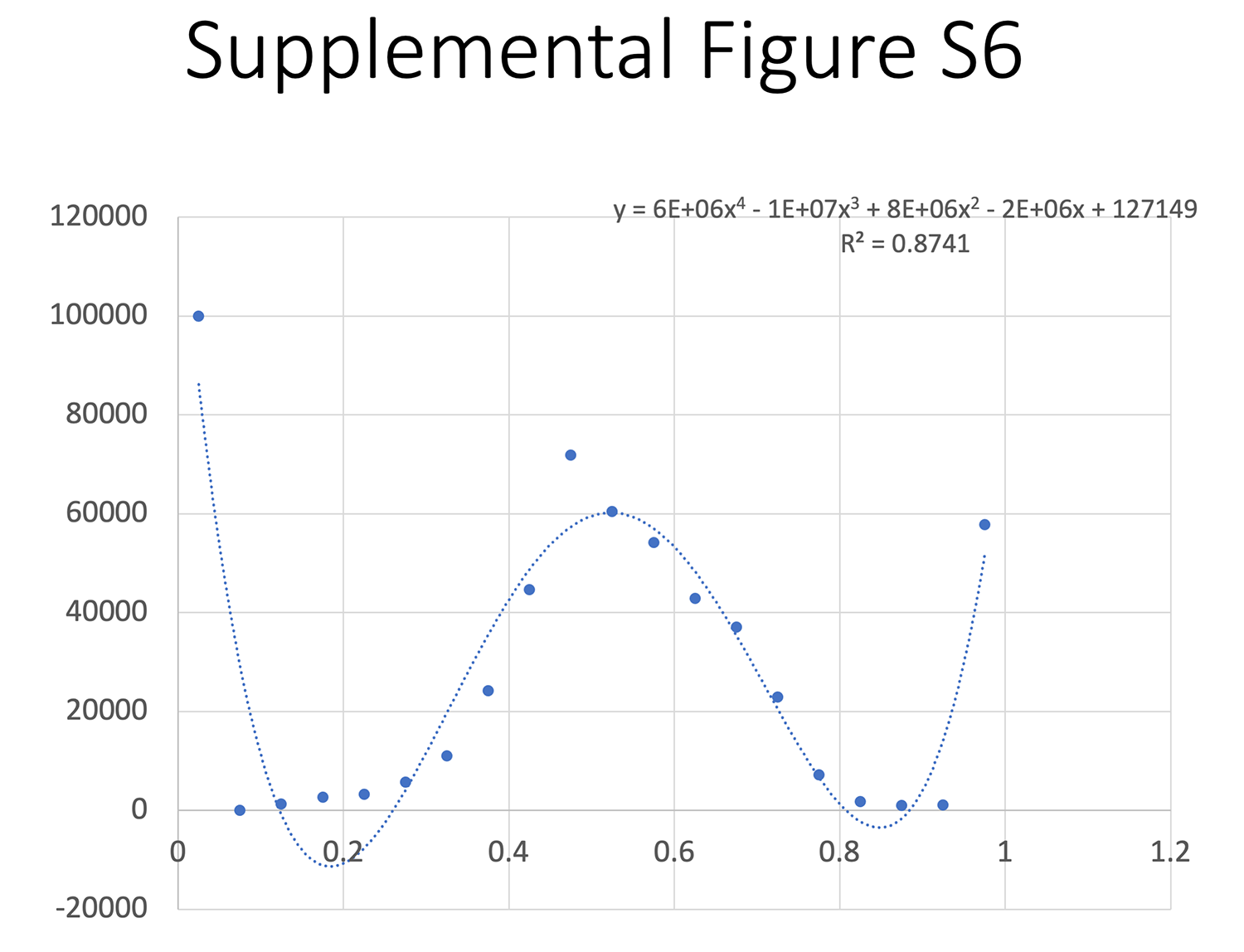

### Supplemental Figure S7

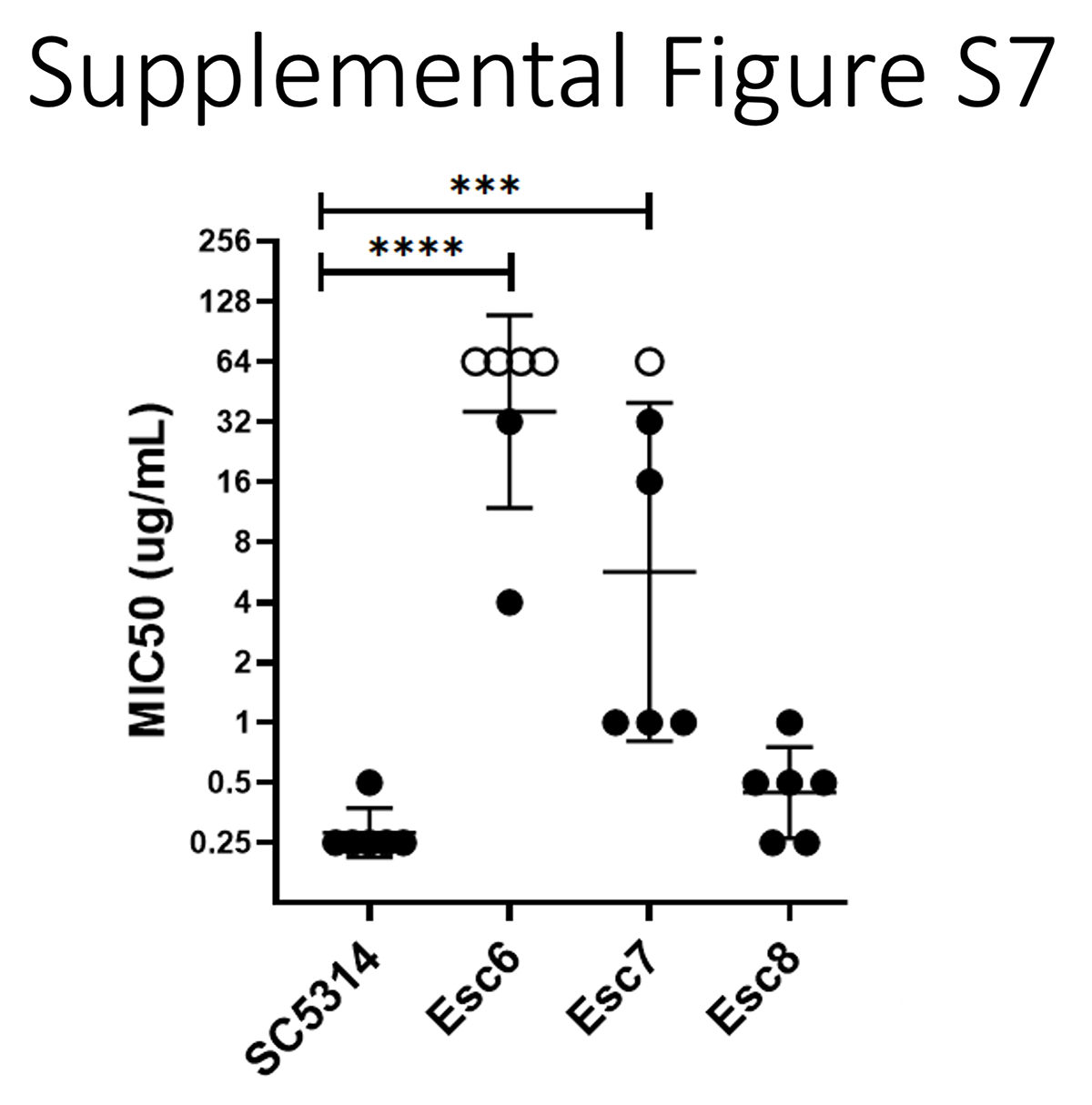

### Supplemental Figure S8

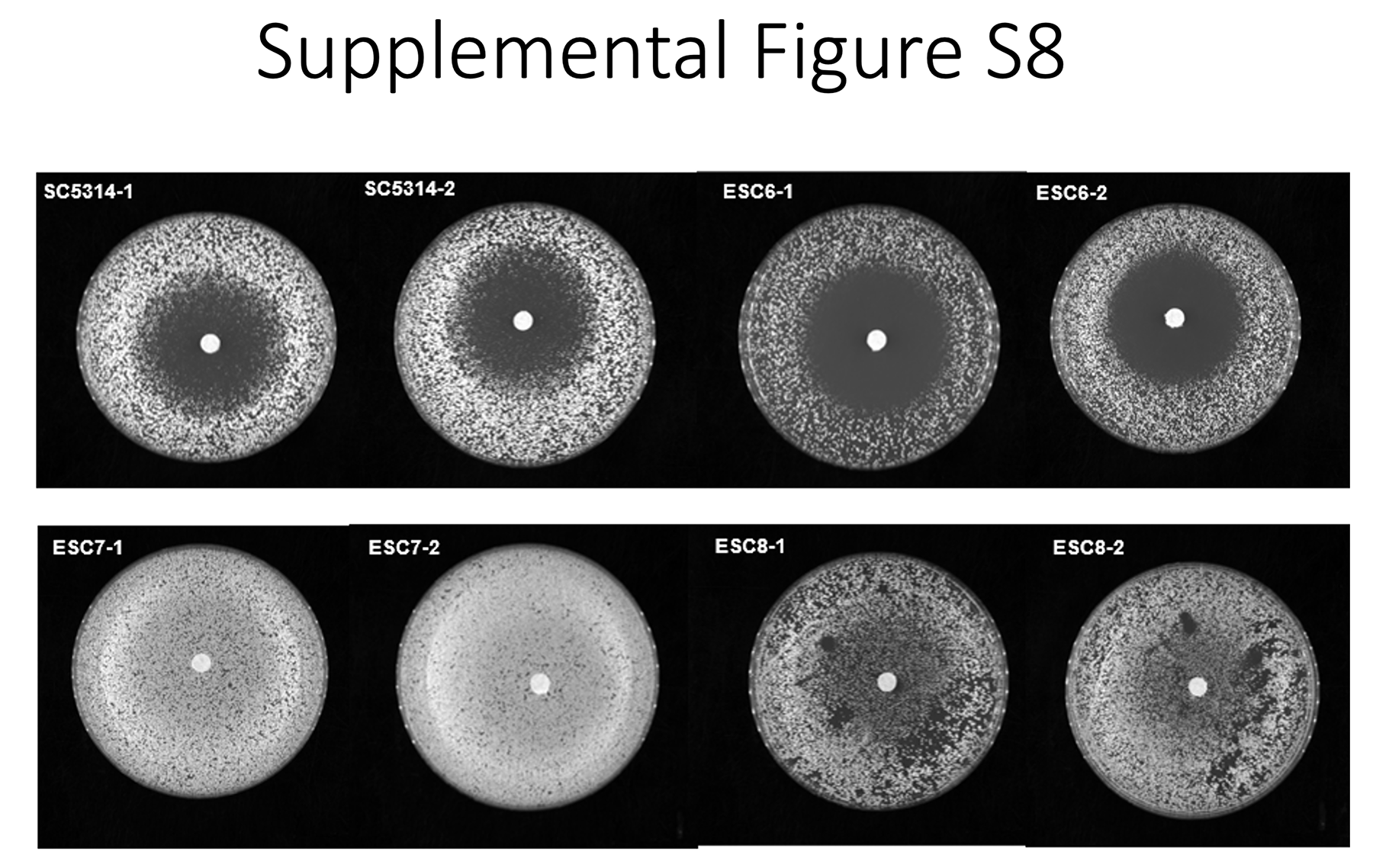

### Supplemental Figure S9

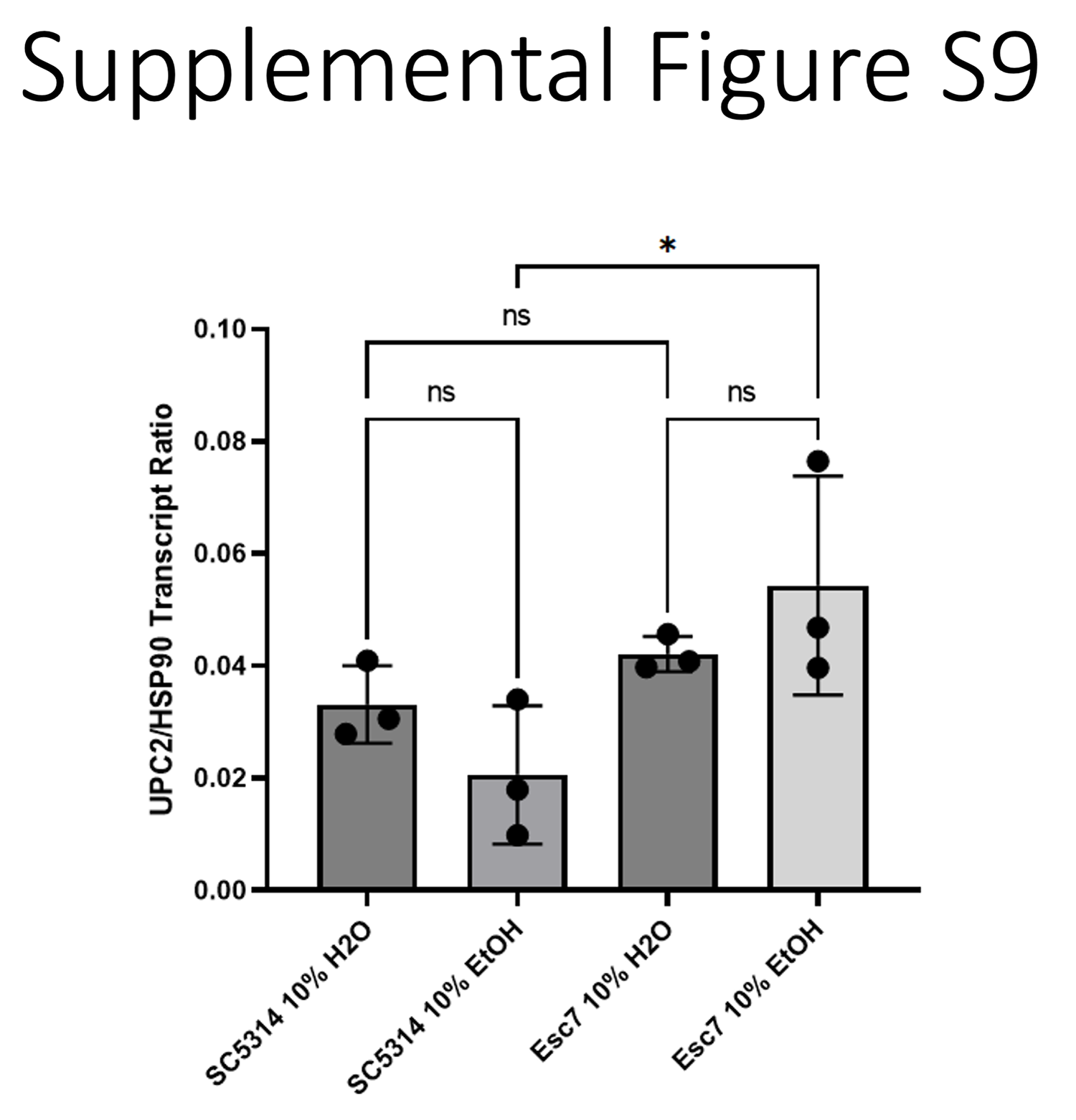
