## Supplemental Table S14 for "Selection of Ethanol Tolerant Strains of *Candida albicans* by Repeated Ethanol Exposure Results in Strains with Reduced Susceptibility to Fluconazole"

Supplemental Table S14 Primers used in this study

| **Primer Name For qPCR^a^** | **Sequence** | **Source** |
| --- | --- | --- |
| ERG11-FW | GCTAATTCTGTTTCATTTAACTCTTCTGAT | [1] |
| ERG11-RV | GGACCAGCTTCGGTATCCAAA |  |
| ERG2-FW | CAGCAATTGGGACTGAAGGT | [2] |
| ERG2-RV | TTCGGGAATCAATGCACCAG |  |
| ERG4-FW | CTTCGGAAGGTCAATCTTGG | [2] |
| ERG4-RV | GTCCAAACACCGGGTAAAGT |  |
| ERG5-FW | GAAGAGCAATTGCGTGTGAG | [2] |
| ERG5-RV | TGGTGGACGGTATCTCAAAG |  |
| ERG6-FW | AGATGCTGCTTCTGTTGCTG | [2] |
| ERG6-RV | GGAATGAAGAACCCCAACC |  |
| ERG25-FW | TATTTCATTGGTGGATACTCTTCATCTT | [1] |
| ERG25-RV | GGACCAGCTTCGGTATCCAAA |  |
| TPS1-FW | TCGCAAGGGTGTCTTGATCTTAT | [3] |
| TPS1-RV | AACAATCAAGGCACCATTAAGTGA |  |
| TPS2-FW | TTGCTGTTGGTCCTGCATCA | [3] |
| TPS2-RV | GGCGAGGTTCGTTCAAATGT |  |
| CDR1-FW | GGTCAACTTGTAATGGGTC | [4] |
| CDR1-RV | AGGACGATAAAGGGCATA |  |
| HSP90-FW | GGGAATCTAACGCTGGTGGTAA | [5] |
| HSP90-RV | TTCGGTTTCTGGAACTTCTTTT |  |
| ADH1-FW | CACTCACGATGGTTCATTCG | [6] |
| ADH1-RV | AAGATGGTGCGACATTGG |  |
| ADH2-FW | AAATGGTTGAACGGCTCTTG | [7] |
| ADH2-RV | GACGGTGACACCAGCACATAAG |  |
| ADH3-FW | ATTCCGACAAATACATTAAAATTAGAGG | This Study |
| ADH3-RV | AATAACCACCAAAATTGAAAATAACTT |  |
| ADH4-FW | TACTGATTCTTATGGATTATATCAAGGA | This Study |
| ADH4-RV | AAAAATCTAACAATTGGAACAATATCAG |  |
| ADH5-FW | ACCTGCAAGGGCTCATTCTG | [8] |
| ADH5-RV | CGGCTCTCAACTTCTCCATA |  |
| ALD4-FW | TTATGCCGTTGAATGTGCTC | [7] |
| ALD4-RV | CTTTGCCCGTGATTTTATCAGC |  |
| ALD5-FW | TGTTGTTACCGGTGGTGCTA | [7] |
| ALD5-RV | CAACGGCTTCGTCAACAGTA |  |
| ACS1-FW | ATTTGCCAGCTTGGTTCATC | [7] |
| ACS1-RV | CACCCTTTTTAACCCCCAAT |  |
| ACS2-FW | CTCAAGGATTTTTCGGTCCA | [7] |
| ACS2-RV | ATTCACCACCCAAAAACCAA |  |
| MDR1-FW | ACATAAATACTTTGCCCATCCAGAA | [9] |
| MDR1-RV | AAGAGTTGGTTTGTAATCGGCTAAA |  |
| SPL1-FW | AAAGGATACTGTGTTAGTTTCTATTATG | This Study |
| SPL1-RV | TATTTTCACGACATATTTTACCAATTTC |  |
| ACT1-FW | GTTGGTGATGAAGCCCAATC | [10] |
| ACT1-RV | CCCAGTTGGAAACAATACCG |  |
| CDR2-FW | GCCAATGCTGAACCGACA | [4] |
| CDR2-RV | ACCAGCCAATACCCCACA |  |

^a^Forward (FW) and reverse (RV) sequences used for primers and their corresponding gene target are listed. For primers designed for this study using NCBI primer blast, the targets were verified by Sanger sequencing.
